# EV*trace*: tracing extracellular vesicles-associated proteins in recipient cells using stable isotope labeling

**DOI:** 10.1101/2025.11.23.690082

**Authors:** Marije E. Kuipers, Joaquin Seras-Franzoso, Mirjam Damen, Richard Wubbolts, Ibane Abasolo, Kelly E. Stecker, Esther N.M. Nolte-‘t Hoen

**Affiliations:** Department of Biomolecular Health Sciences, Division of Infectious Diseases and Immunology, Faculty of Veterinary Medicine, Utrecht University, Utrecht, The Netherlands; Clinical Biochemistry, Drug Delivery and Therapy Group (CB-DDT), Vall d’Hebron Research Institute (VHIR), Vall d’Hebron University Hospital, Barcelona, Spain; Centro de Investigación Biomédica en Red, Bioingeniería, Biomateriales y Nanomedicina (CIBER-BBN), Barcelona, Spain; Biomolecular Mass Spectrometry and Proteomics, Center for Biomolecular Research and Utrecht Institute for Pharmaceutical Sciences, Utrecht University, The Netherlands; Center for Cell Imaging, Department of Biomolecular Health Sciences, Faculty of Veterinary Medicine, Utrecht University, Utrecht, The Netherlands; Institute for Advanced Chemistry of Catalonia (IQAC-CSIC), Barcelona, Spain

## Abstract

Release and uptake of extracellular vesicles (EVs) is a highly conserved means of communication used by cells across all kingdoms of life. These nano-sized vesicles transfer messages encoded in proteins and nucleic acids and play a role in numerous (patho)physiological processes. A longstanding question in the field is whether interaction and uptake of EVs is a stochastic process or whether specific EV subpopulations target different types of recipient cells. Here we present *EVtrace*, a proteomics-based approach that accommodates stable isotopic labeling of amino acids in culture (SILAC) to trace back labeled EV-associated proteins in unlabeled recipient cell types. We describe the optimization of EV labeling conditions and EV-cell cultures, and introduce a proteomics data analysis pipeline to confidently identify sparse internalized EV proteins among unlabeled recipient cell proteins. As a proof of concept, we studied the targeting of prostate cancer (PCa) EVs to bone cells and non-bone cells, considering that bone is a common metastatic site of PCa. Using *EVtrace* we demonstrate that interaction of PCa EVs with recipient cell types was not stochastic, and that different subpopulations of EVs targeted different cells. Proteins of EV subpopulations that were traced in bone cells, but not in non-bone cells, were strongly enriched for pathways involved in cancer progression and metastasis. With super-resolution microscopy we confirmed that EV proteins traced in different targeted cell types co-occurred on specific EV subpopulations. EV*trace* is a valuable new tool for EV research because it supports identification of EV subpopulations that are transferred to recipient cells and discloses candidate proteins potentially involved in EV binding/uptake and functional modification of target cells.

## Introduction

Extracellular vesicles (EVs) are part of the signaling network between cells^1^. The term EVs includes all particles that are released by cells into the extracellular space that are delimited by a lipid bilayer and cannot self-replicate^2^. Cells can release several EV subtypes via different routes of biogenesis and release^3^. Exosomes are formed as intraluminal vesicles by inward budding of intracellular compartments, such as endosomes or autophagosomes, and can be released as EVs upon fusion of these compartments with the plasma membrane. These EVs have a size range of ∼30-150 nm. EVs with a larger size range (∼50-10,000 nm) can be formed by direct outward budding of the plasma membrane or when cells undergo apoptosis. Each of the biogenesis mechanisms involves a different set of proteins and lipids, which causes heterogeneity in the molecular composition of EVs released by one cell type. This heterogeneity is further increased by variation in the activation- and differentiation-status of EV-producing cells. Depending on their molecular composition, EVs can modify the function of receiving cells, provide trophic support, or modulate the extracellular matrix^3^. Related to their diverse functions, EVs have been implicated in various pathologies, and are explored for their biomarker and therapeutic potential^4^.

Profound understanding of the biological function of EVs requires fundamental understanding of how EVs interact with their recipient cells, through surface binding and/or internalization^5^. However, despite our increasing knowledge on EV heterogeneity, it remains unclear how diversity in the molecular composition of EV subsets relates to differences in their interaction with target cells and function. EV-binding and -uptake is often assumed to be a stochastic process, whereby a random fraction of the total EV population is internalized. This assumption, however, is based on experiments in which bulk-isolated EVs labeled with generic luminal or membrane dyes are added to recipient cells^6,7^. This methodology does not provide information on EV subpopulation-specific differences in binding and uptake by target cells. Beyond uptake efficiency, little is known regarding the type of molecules that determine whether EVs are internalized and functionally active within the EV-recipient cell.

Omics-based analyses of bulk EVs, such as proteomics, can identify thousands of EV-associated molecules. However, these approaches do not resolve which EV-associated molecules are transferred to the cells, nor do they capture the combinatorial diversity of molecules within individual EVs. Targeted strategies using (fluorescent) tagging or antibody labeling yield knowledge on specific EV subpopulations^8^, but are limited to the investigation of preselected proteins or lipids. Furthermore, molecule tagging that requires cell engineering^7^ can alter EV composition relative to native EVs. The unresolved questions regarding target cell specificity, uptake efficiency, and the molecular composition of internalized EV subpopulations, point to a critical gap in our understanding of how EV composition relates to their biological impact. There is an urgent need for unbiased technologies that can track multiple EV-associated molecules simultaneously and correlate them with cellular uptake and downstream function. Such approaches are essential to uncover how the molecular composition of EV subpopulations shapes their interactions with specific target cells and ultimately governs their biological roles.

To address the current knowledge gap, we developed a novel SILAC (stable isotope labeling by amino acids in cell culture) proteomics-based method which we coined ‘EV*trace*’. This method allows unbiased tracing of stable isotope-labeled EV proteins in EV-recipient cells.

Standard SILAC-based protein quantification by mass spectrometry (MS) is performed by ratiometric comparison between labeled and unlabeled versions of the same proteins in multiple conditions. This approach has been applied in EV research to study changes in protein composition of EVs between different EV isolations^9^, culture conditions^10–12^ or cell types^13,14^. Thus far, SILAC approaches have only been used for targeted analysis of pre-selected EV proteins without considering the comparison to the full EV proteome^15–18^. Our novel EV*trace* method is a SILAC proteomics approach for unbiased tracing of labeled EV proteins in unlabeled EV-recipient cells. We here describe the development of the tracing method and the accompanying proteomics data analysis pipeline to confidently identify sparse internalized EV proteins among the abundant unlabeled recipient cell proteins. To obtain proof of concept for EV*trace*, we utilized prostate cancer (PCa) EVs known to target bone cells, particularly osteoblasts and osteoclasts^19,20^. We investigated the number and identity of EV proteins that could be traced back in different bone cells types, monocytes, and tumor cells. Results showed that the EV protein profiles traced in the four cell types differed significantly from one another, suggesting specific targeting of EV (sub)populations to different recipient cell types. Comparison of the traced EV proteins to the abundance of these proteins in the source EV pool indicated that detection of EV proteins on/inside cells is not a simple reflection of their overall abundance. Furthermore, enrichment analyses showed that the EV proteins traced back in bone cells displayed an increased representation in biological processes associated with cancer. These findings demonstrate that EV*trace* can be used as an unbiased method to identify EV proteins among a total pool of EV proteins that are prone to transfer to EV-recipient cells of interest. EV*trace* generates valuable sources of candidate proteins that can be further explored for their role in EV binding/uptake and functional modification of target cells.

## Methods

### Cell cultures

#### EV-donor cell cultures

PC3 (RRID:CVCL_0035) cells (bought from ATCC, Manassas, VA, USA) were cultured in RPMI 1640 Glutamax (Gibco, Thermo Fisher Scientific, Waltham, MA, USA) supplemented with 10% FBS (Serana Europe GmbH, Pessin, Brandenburg, Germany) and Penicillin-Streptomycin (pen/strep) (Gibco) in T75 flasks (Corning Inc., Corning, NY, USA) at 37 °C and 5% CO_2_. They were passed twice a week using 0.05% trypsin (Gibco) and not further than passage number 25. The cell cultures were frequently tested for the absence of mycoplasma. For the SILAC cultures, PC3 cells from regular cultures were diluted 1:10 into SILAC RPMI 1640 Flex media (Gibco) supplemented with 10% dialyzed FBS (Gibco), 2 mM Glutamax (Gibco), 0.1 mg/ml L-Arganine:HCL (13C6;15N4) (Cambridge Isotope Laboratories, Andover, MA, USA), 0.04 mg/ml L-Lysine:2HCL (3,3,4,4,5,5,6,6-D8) (Cambridge Isotope Laboratories), 0.2 mg/ml L-Proline (Sigma-Aldrich, St. Louis, MO, USA) and pen/strep. Cells were passed at least 4 times in SILAC medium before seeding for EV collection.

For collection of EVs, PC3 cells were cultured in T175 flasks (Greiner Bio-One International GmbH, Kremsmünster, Austria) till 85% confluency. The medium was removed and cells were washed with PBS (Gibco) before new culture medium was added with 10% EV-depleted (dialyzed) FBS and the supplements as described above. The 10% EV-depleted FBS was made from a stock of 30% EV-depleted FBS in RPMI (SILAC or regular). For this, 30% FBS was centrifuged for >16 h in polypropylene tubes at 28,000 rpm and 4 °C in an SW32 Ti rotor and L90K ultracentrifuge (Beckman Coulter Life Sciences, Brea, CA, USA). Subsequently, the top 25 ml of each tube was collected, filtered (0.22 µm filter, Corning), aliquoted, and stored at −20 °C. Culture medium was collected from the cells after 16-18 h, and subjected to two times 200 × *g* and two times 500 × *g* centrifugation steps. Each centrifugation step was 10 min and at 4 °C, after which the supernatant was decanted to a new 50 ml tubes before the next centrifugation step. The final 500 × *g* supernatant was transferred to new tubes using a serological pipette, aliquoted, and stored at -80 °C till EV isolation.

#### EV-recipient cells cultures

PC3 cells were cultured in RPMI as described above. SAOS2 (RRID:CVCL_0548) cells (kind gift from Prof J.H.E. Kuball, University Medical Center Utrecht) were maintained in McCoy 5a Glutamax medium (Gibco) supplemented with 15% FBS and pen/strep. THP1 (RRID:CVCL_0006) cells (ATCC) were maintained in similar medium as for the PC3 cells with the addition of 0.05 mM β-Mercaptoethanol. Cells passaged less than 20 times were used for experiments. Cells were seeded in 48 well plates (Corning) at various concentrations (20,000-35,000 cells per well for PC3 and SAOS2, 40,000-60,000 cells for THP1) 24 h before EV incubation. For the calculation of protein concentration related to cell numbers, 24 h after seeding, cells were washed with PBS and lysed for 30 min on ice in RIPA buffer (40 mM Tris-HCl pH 8, 1% Triton X-100, 0.5% sodium deoxycholate, 150 mM sodium chloride, 0.1% sodium dodecyl sulfate) with the addition of proteinase inhibitor (Roche, Basel, Switzerland). Lysed cells were centrifuged at max. speed in an Eppendorf centrifuge for 15 min and the protein concentration was calculated in the supernatant by a BCA protein assay (Pierce, Thermo Fisher Scientific) following the manufacturers protocol. RIPA was used as the BCA background.

Osteoclasts were differentiated from THP1 cells as described elsewhere^21^. THP1 cells were harvested, pelleted by centrifugation at 150 × *g* and resuspended in full THP1 RPMI culture medium without β-Mercaptoethanol. Cells were seeded in flat bottom 48 well plates (50,000-100,000 cells/well) in the presence of 100 ng/ml PMA (Sigma-Aldrich). PMA differentiates the THP1 cells first to macrophage-like cells and also stops the cells from dividing. After 48 h, half of the medium was replaced with RPMI culture medium supplemented with 10% FBS and pen/strep. After 24-72 h of rest, cells were differentiated to osteoclasts by addition of 30 ng/ml human M-CSF (GenScript Biotech, Rijswijk, the Netherlands) and 50 ng/ml recombinant human RANK-L/TRANCE (R&D Systems, Minneapolis, MN, USA), which was refreshed every 2 days. After 7-9 days in differentiation medium, the osteoclast were ready to be incubated with EVs. Differentiation to multinuclear osteoclasts-like cells was confirmed by TRAP staining following the manufacturers protocol (Sigma-Aldrich) as well as CD51 and CTSK immunolabeling and CTSK activity assay (Supplementary Figure 1). An equivalent THP1 differentiation protocol was employed for determination of CTSK enzymatic activity and flow cytometry characterization, which were performed using initial amounts of 175,000 THP-1 cells/well in flat bottom 24 well plates.

#### CTSK enzymatic activity assay

THP-1 cells, macrophages and osteoclasts were lysed in 50 mM MES buffer (pH 5.5 and 1% (v/v) Triton X-100) on ice for 20 min. A cell scraper was used to improve sample recovery. Samples were subsequently sonicated twice for 20 s at 60% amplitude, 0.5 s on/off. Total protein concentration was inferred by BCA assay according to the manufacturer’s instructions. Cell lysates were diluted in reaction buffer (50 mM MES, 2.5 mM EDTA, 2.5 mM DTT, 10% DMSO) at final concentration of 0.1 mg total protein/ml and 100 µl of sample was dispensed into 96 well black plates. CTSK activity was monitored by measuring the cleavage of Z-Leu-Arg-7-amido-4 methylcoumarin (Sigma-Aldrich, 164545) substrate peptide, rendering a fluorescent sub product (AMC). 100 µl of either CTSK substrate at 10 µM, or substrate plus CTSK inhibitor (44 nM, Sigma-Aldrich, 219377) were added to the samples and incubated 1 h at RT protected from light. Fluorescence was measured in a Varioskan LUX plate reader (Thermo Fisher) at 375 nm excitation and 460 nm emission.

#### Cell immunolabeling for flow cytometry

Cells were detached with accutase and washed in PBS/BSA (0.5 % w/v), counted, and 500,000 cells aliquoted per condition in 1.5 ml tubes. 5 µl of CD51-PE antibody (Biolegend 327909, clone NKI-M9) was added per sample in PBS/BSA, 100 µl final volume, and incubated on ice for 30 min with gentle agitation in the dark. After incubation, cells were pelleted and washed in 1 ml PBS/BSA. Samples were subsequently fixed in 2% PFA for 15 min at RT, washed with PBS/BSA, and permeabilized in 0.1% (v/v) Triton X-100 in PBS. Samples were blocked for 15 min at RT using human IgG solution at 100 µg/ml after another washing step. Cells were then pelleted and incubated with 10µg/ml CTSK polyclonal primary antibody (Invitrogen PA5-14270) for 1 h on ice. An additional washing step was carried out before 20 min incubation on ice with the secondary AF488 conjugated antibody (Invitrogen A-21441) at 2 µg/ml. Finally, cells were washed in PBS/BSA and resuspended in PBS. Samples were analyzed in a Fortessa flow cytometer (BD Biosciences). Gates were set based on unstained and single stained controls.

### EV isolation

The preparation of one EV batch for cell stimulation and super-resolution microscopy started from 120-210 ml of cell culture medium and EV batches for western blot and high-resolution flow cytometry were from 20-60 ml culture medium. All ultracentrifugation tubes were open-top thinwall polypropylene, all centrifugation steps were done at 4 °C, and the ultracentrifuges used were an L90K and an Optima XPN-80 (Beckman Coulter). The BSA used was from a 5% BSA in PBS stock that was depleted of aggregates by ultracentrifugation as previously described^22^. We have submitted all relevant data of our experiments to the EV-TRACK knowledgebase (EV-TRACK ID: EV250121)^23^.

500 × *g* culture supernatants were thawed overnight at 4 °C, transferred to SW32 tubes, and centrifuged for 30 min at 8,900 rpm (average of 9,759 × *g*, k-factor of 2,637) in an SW32 Ti rotor. The supernatant (referred to as 10,000 × *g* supernatant) was transferred to new SW32 tubes and the pellet was resuspended in 100 µL PBS+0.2% BSA for density gradient separation or discarded. The 10,000 × *g* supernatant was further centrifuged for 65 min at 28,000 rpm (average of 96,589 × *g*, k-factor of 266) in an SW32 Ti rotor. The supernatant (referred to as 100,000 × *g* supernatant) was removed down to the conical part of the tubes, after which the 100,000 × *g* pellets were resuspended in the remaining supernatant, transferred and pooled into two SW40 tubes, topped up with 1-3 ml 100,000 × *g* supernatant, and centrifuged for 65 min in an SW40 Ti rotor at 28,000 rpm (average of 99,223 × *g*, k-factor of 279). Subsequently, the supernatant was removed till the conical part of the tube after which the tube was carefully decanted to remove the remaining liquid. The walls of the tube were dried with a tissue while the tube was still upside down. The EV-enriched pellet was resuspended in 80-200 µl (SILAC labeled EVs for EV*trace*, western blot, and super resolution microscopy) or 40 µl (for EV fluorescent labeling) PBS+0.2% BSA. EVs were labeled with PKH26 by adding 60 µl of diluent C to the resuspended EV-enriched pellet followed by a 3 min incubation with 93 µl PKH26 (1,5 µL diluted in 100 µL diluent C) (Sigma-Aldrich) after which the labeling was quenched by addition of 100 µL RPMI+10% EV-depleted FBS. RPMI+10% EV-depleted FBS was processed and labeled with PKH26 similar to the EVs as a dye control. Resuspended EV pellets were transferred to SW60 tubes and gently mixed with 970 µL 60% iodixanol (Optiprep, Stemcell Technologies, Vancouver, Canada). On top of this, 485 µl 40%, 485 µl 30%, and 1,746 µl 10% iodixanol layers were added. These iodixanol dilutions were prepared from a 50% dilution and PBS (Gibco). The 50% iodixanol was made from 60%, 10x PBS (Gibco), and sterile MilliQ (MQ) water. All density gradients were centrifuged 16-18 h in an SW60 Ti rotor at 43,000 rpm (average of 191,556 × *g*, k-factor of 87). From the density gradients, 12 equal fractions were collected from top to bottom (where the top fraction was named F12 and the bottom fraction F1). Fractions were analyzed by flow cytometry and their density measured in an Abbe refractometer (Atago, Tokyo, Japan). Iodixanol density of each fraction was calculated from the refractive index (RI) by the formula (3.35*(RI))-3.4665. The fractions for the western blots were subsequently subjected to a TCA protein precipitation protocol as described previously^24^. For all cell incubation experiments (with SILAC EVs or PKH26-labeled EVs) and super resolution microscopy, fractions 6 and 7 (with iodixanol densities between 1.10-1.06 g/ml) were pooled and directly mixed with 11 ml of PBS+0.1% BSA in SW40 tubes. The tubes were centrifuged for 65 min in an SW40 Ti rotor at 39,000 rpm (average of 192,498 × *g*, k-factor of 144) after which the supernatant was removed and tube was dried upside down as mentioned above. The final isolated EV pellets were resuspended in 40 or 70 µl PBS+0.2% BSA for PKH-labeled EVs and SILAC EVs, respectively. EV pellets for super-resolution microscopy were resuspended in 66-88 µl of PBS+0.2% BSA, aliquoted, and stored in -80 °C. For the resuspended SILAC EVs, an aliquot of ∼30 µl was directly mixed with 5× reducing Laemmli sample buffer, heated 5 min at 98 °C, and stored at −20 °C. All other purified EVs were directly added to cells after isolation.

### Nanoparticle tracking analysis

EV aliquots from cell experiments (stored at 4 °C) were measured on the same day by nanoparticle tracking analysis (NTA) as described elsewhere^25^. Briefly, EVs were diluted 500× in PBS and three recordings of 30 s were made on camera level 16, 14, and 12 each. A NanoSight NS500 (Malvern Panalytical, Malvern, UK) equipped with an sCMOS camera was used, and data was acquired and analyzed in NTA3.3 software with a detection threshold of 5. PBS alone was measured in similar manner and used for background subtraction. Uncultured RPMI + 10% EV-depleted FBS was processed and measured similar to the EVs as medium control.

### Western blot

Trichloroacetic acid (TCA) precipitated proteins from density gradient fractions were resuspended in non-reducing (CD63 and CD9) or reducing (TSG101) Laemmli sample buffer, heated for 5 min at 98 °C, and stored at −20 °C till loading on gel. Samples (15 µl/lane) and marker (Precision Plus Protein Dual Color, Bio-Rad Laboratories, Hercules CA, USA) were run through a 12.5% SDS-PAGE gel and subsequently transferred to a PVDF membrane (0.2 µm, Bio-Rad). Membranes were blocked with PBS supplemented with 0.2% gelatin from fish skin and 0.1% Tween20 and incubated overnight with antibodies: CD63 (1:500, clone TS63, 106280, Invitrogen, Thermo Fisher Scientific), CD9 (1:2,000, clone HI9a, 312102, BioLegend, San Diego, CA, USA), TSG101 (1:1,000, rabbit polyclonal, T59512, Sigma-Aldrich). Blots were washed 5 times with blocking buffer and subsequently incubated with goat-anti mouse IgG+IgM (H+L)-HRP (1:10,000, polyclonal, 115-035-044, Jackson ImmunoResearch, West Grove, PA, USA) or goat-anti rabbit Ig-HRP (1:5,000, polyclonal, P044801-2, Agilent Technologies, Santa Clara, CA, USA) for 45 min on a rocking platform. Blots were washed 5 times with blocking buffer, 3 times with PBS+0.1% Tween20, and 3 times with PBS before ECL detection (SuperSignal West Dura, Thermo Fisher Scientific) in a ChemiDoc imager (Bio-Rad).

### High-resolution flow cytometric analysis of EVs

To observe the distribution of EVs among the density gradient, gradient fractions from PKH26-labeled PC3 EVs were measured on an Aurora spectral flow cytometer with small particle detector (Cytek Biosciences, Fremont, CA, USA). PBS alone and isolated unstained EVs were used as (background) controls to set the threshold on the B4 detector (highest peak detector for PKH26 scatter), allowing only PKH26-labeled EVs above the threshold to be recorded (Supplementary Figure 2). This B4 threshold was set on 1,100. The gains of the detectors were manually set to the following: V1/V2/V11/V12/FSC/SSC = 2,000; V3/B4 = 1,250; V4/R8 = 750; V5-V10/V13-V16/B6-B14/R1/R3-R7 = 3,000; B1-B3/R2 = 2,500; B5 = 1,500. Flow rate was set on low and 20 µl of each sample that was diluted 50× in PBS (Gibco) was recorded. The analysis was done in FlowJo (version 10, Becton Dickinson(BD) Biosciences, Franklin Lakes, NJ, USA).

### Super-resolution microscopy

Density gradient-purified PC3 EV aliquots were thawed and diluted 1:1 in PBS+0.1%BSA. EVs were prepared using the EV Profiler and Profiler 2 Kits (ONI, Oxford, UK) according to the manufacturers’ instructions, with the exception of leaving out the fixation step after EV capture. The EV capturing method was phospholipids-based, and all steps of the capturing and labeling were performed at RT. Antibodies CD9-CF488A, CD63-CF568 and CD81-CF647 were provided in the EV profiler kit. Anti-GRP78 (clone C38, *InVivo*MAb, Bio X Cell, Lebanon, NH, USA) was labeled by anti-mouse IgG2a/b-VHH (N2705, NanoTag Biotechnologies, Göttingen, Germany) that was labeled in house with AlexaFluor647. Unconjugated integrin-β1 antibody (clone P5D2, Santa Cruz Biotechnology, Dallas, TX, USA), CD47 antibody (clone B6.H12, *InVivo*MAb), or mouse IgG1 isotype control (clone P3.6.2.8.1, eBioscience, Invitrogen) were labeled with mouse IgG1 targeting nanobodies (VHH-AlexaFluor568, VHH-AlexaFluor647, Proteintech, Rosemont, IL, USA). For this, 3 µg of the VHH was incubated with 5 µg of antibody for 1 hour at 4 °C. Unbound VHH was washed away by spinning the mixture 15 min on 12,000 ×*g* in 50k MWCO PES Vivaspin filters (Sartorius, Göttingen, Germany). All antibodies were centrifuged briefly before use to avoid aggregates. All antibodies used for EV staining had the final concentration of 7.5 µg/ml, and staining was done for 50 min. The WGA (Wheat Germ Agglutinin) staining (5 min, 1:500 of 2.5 mg/ml, Invitrogen) was done after sample fixation, just before adding the dSTORM buffer provided in the EV Profiler Kit (ONI). Image acquisition was done on a Nanoimager (ONI). For each sample, 3 fields were acquired with the following protocol: 6,000-10,000 frames 640 (30% laser power), 6,000-10,000 frames 561 (50% laser power), 6,000-10,000 frames 488 (50% laser power), with 30 ms exposure time. The first 300-500 frames of each laser were discarded in the analysis. The analysis was done in CODI (ONI) using the EVprofiling settings. For the tetraspanin staining in Figure 1, DBSCAN clustering was done using a min sample size of 10 and min size of 20 and 70 nm distance and the circularity was set to 0.8-1. For the other staining panels, filters were adjusted from the EVprofiling settings: photon count 200-100,000, sigma 50-300 nm, localization precision 0-15 nm, and background photons 1,500-200,00. DBSCAN clustering was done by merging the two channels and using a min sample size of 5 and min size of 10 and 70 nm distance. The filtering of the clusters was set to an area of min 1000 nm^2^, circularity of 0.7-1, 20-1,000 nm radius of gyration, and 0.001-1 per nm^2^ density. The counting was based on 10 rings, a bin size of 20, max r value of 300, with a count threshold of 2, and the counting radius was set at 120 nm.

**Figure 1.**
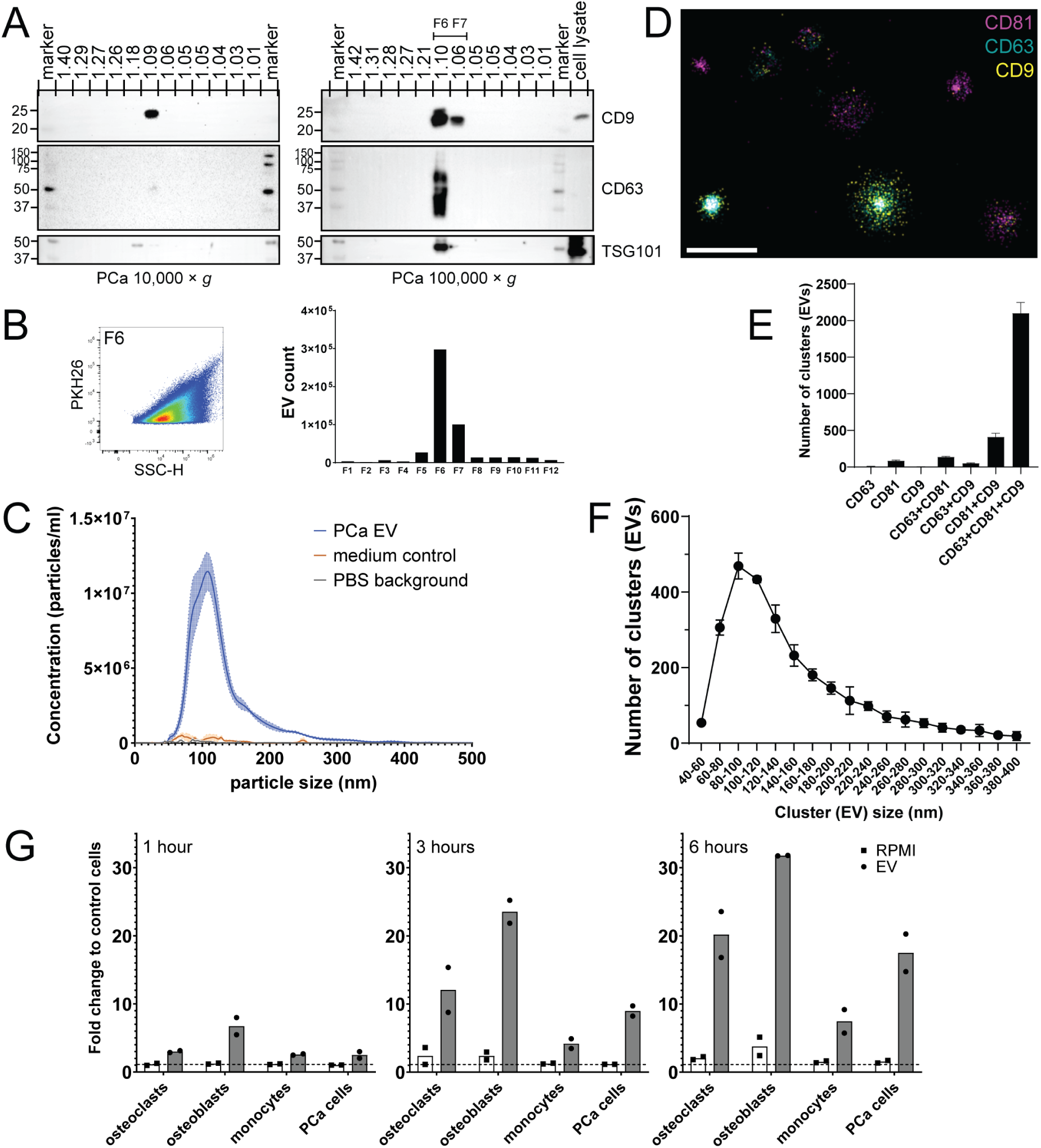
Characterization of isolated PCa EVs and interaction with EV-recipient cells. (**A**) Western blot detection of EV-enriched proteins CD9, CD63, and TSG101 in iodixanol density gradient fractions of 10,000 × *g* and 100,000 × *g* PCa EV pellets. Fraction densities are indicated as g/ml. Data are representative of 2-5 independent experiments. (**B**) Fluorescently labeled EVs (PKH26) in each density fraction (as in A) were measured by high-resolution flow cytometry. Indicated is an example dotplot of Fraction 6 (left) and EV counts per fraction (right). (**C**) NTA of EVs isolated from F6+F7 density fractions in A (blue), medium control (orange) and PBS (grey). Data are presented as mean values ±SEM of 2 individual EV isolations measured on 3 different camera levels. (**D-F**) Super-resolution microscopy of PCa EVs stained for CD81, CD63 and CD9. (**D**) Close up image of multiple clusters (EVs) stained for the indicated tetraspanins (size bar indicates 500 nm). (**E**) Quantification of the co-localization of various tetraspanins on each measured cluster of three fields of view. A cluster is indicative of 1 EV and data are presented as mean values from 3 fields of view ±SEM. (**F**) Size distribution of clusters (EVs). Average data from 3 fields of view ±SD. (**G**) The indicated cell types were incubated with fluorescently labeled PCa EVs (PKH26) (grey bars) or dye control (white bars) for 1, 3 or 6 h and measured by flow cytometry. The fold change indicates the amount of EV interaction with the cells compared to control cells that were not incubated with EVs or dye control. PCa, prostate cancer

### EV-cell incubations

One hour before EV incubation, light microscopic images were acquired of the cells in the wells (5 images of 1 well of each cell concentration) and surface area coverage was calculated per cell type in ImageJ Fiji^26^ using the PHANTAST plugin^27^ and in house generated macro scripts (Supplementary Figure 3A). Briefly, noise removal, median filter, CLAHE, and background removal were applied before running the PHANTAST plugin. PHANTAST settings used were epsilon 0.03 and sigma 0.6 (THP1), 0.7 (osteoclasts) or 1.7 (PC3 and SAOS2 cells). Subsequently, post segmentation closing and erosion (3 px) were applied before binary area calculations. The average of the images per cell concentration were used to select the wells with osteoblasts, monocytes, and PCa cells with surface coverages that were closest to the coverage of the wells with osteoclasts. Similar numbers of EVs per calculated cell surface coverage were added to each cell type and volumes in the wells were adjusted to keep EV concentrations the same between all cell types. For the PKH26-labeled EVs, the highest amount of EVs added (to the cells with the most surface coverage) was from half a T175 flask (equal to 15 ml of EV collection medium). An equal volume of the PKH dye control sample was added to the cells. For the EV*trace* experiments, the maximum amount of added EVs was derived from one T175 flask, which was equal to 30 mL of EV collection medium.

After 1, 3 and 6 h of incubation of the cells with the PKH26-labeled EVs, cells were washed with PBS and harvested after accutase-based cell dissociation (Sigma-Aldrich) before flow cytometric measurement (Canto II, BD Biosciences). Data was analyzed in FlowJo (version 10). EV-cell interaction between the different cell types was determined by calculating the fold change of the sum of mean fluorescent signals of equal numbers of cells with and without EV incubation. This allowed the inclusion of all EV interactions, independent of whether it was a few cells that interacted with many EVs or whether many cells interacted with only a few EVs. Graphs of flow cytometry data were made in GraphPad Prism (version 10, Dotmatics, Boston, MA, USA).

After 3 h of incubation of the cells with the SILAC EVs, osteoclasts, SAOS2, and PC3 cells were carefully washed with PBS and lysed in the plate for 30 min on ice in 30 µl RIPA buffer with the addition of proteinase inhibitor (Roche, Basel, Switzerland). THP1 cells were first transferred to an Eppendorf tube to be washed with PBS by pelleting the cells by centrifugation at 150 × *g* for 5 min. After washing the cells, the pellet of THP1 cells was resuspended in 30 µl of RIPA buffer + proteinase inhibitor, and incubated on ice for 30 min. Lysed cells from the plates were transferred to Eppendorf tubes and all Eppendorf tubes were centrifuged for 15 min at >16,000 × *g* (4 °C). Finally, the supernatants of the cell lysates were transferred to new Eppendorf tubes, mixed with 5× reducing Laemmli sample buffer, incubated for 5 min at 98 °C, and stored at −20 °C.

### Proteomics

#### Sample preparation

Lysed cells and EVs in sample buffer were run through a 4-12% Bis-Tris gel (Criterion XT, 12+2 well, Bio-Rad), after which the gel was fixed (40% Methanol + 10% Acetic Acid in MQ) for 30 min. Gel was washed several times with MQ before 1 hour mixing in protein staining solution (Imperial Protein Stain, Thermo Fisher Scientific). Gel was destained overnight in MQ and imaged on an Amersham Imager 600. Gel fractions were cut as pictured in Supplementary Figure 4, and each fraction was cut into small pieces and transferred to LoBind Eppendorf tubes for in-gel digestion. For this, gels were washed in MQ, followed by double incubation in 100% acetonitrile (ACN) until gel pieces were white. Gel pieces were incubated for 1 hour in 6.5 mM DTT (in ammonium bicarbonate, 50 mM and pH 8.5) followed by two ACN treatments. Next, gel pieces were incubated in 54 mM iodoacetamide (in ammonium bicarbonate) for 30 min in the dark. After two subsequent ACN incubations, gel pieces were washed twice with ammonium bicarbonate, where each wash step was followed by two ACN incubations. Trypsin (Promega, Madison, WI, USA) was resuspended in cold ammonium bicarbonate to a dilution of 3 ng/µl and 10 µl was added to the gel pieces in each Eppendorf tube and incubated on ice for >30 min after which 30 µL ammonium bicarbonate was added each 30 min until all gel pieces were covered. Samples were incubated overnight at 37 °C for digestion. ACN was added to each Eppendorf tube to extract all peptides, and the peptide containing supernatant was transferred to a new Eppendorf tube and dried. Dried peptides were stored at −20 °C till MS measurements.

#### Mass spectrometer measurements

Dried peptides were reconstituted in 10 µl 10% formic acid (FA) and analyzed on an Orbitrap Exploris 480 mass spectrometer (Thermo Fisher Scientific) connected to a UHPLC 3000 system (Thermo Fisher Scientific). Peptides were trapped on a double-fritted precolumn (2 cm x 100 µm, Reprosil C18) (3µm) and then separated on a 50 cm x 75 µm Poroshell EC-C18 analytical column (2.4µm). Solvent A consisted of 0.1% FA, solvent B of 0.1 % FA in 80% ACN. Trapping was performed for 1 min in 9% solvent A and peptides were subsequently separated by a 43 min gradient of 13–55% solvent B followed by 2 min 55–99% solvent B and 5 min 99% solvent B. Flow rate was set to 300 nl/min and the column was heated to 40°C. MS data was obtained in data-dependent acquisition mode. Full scans were acquired in the m/z range of 375-1600 at the resolution of 60,000 (m/z 400) with AGC target 3e6. Precursor ions were continuously selected for HCD fragmentation over the course of 2 s. Fragmentation was performed at normalized collision energy (NCE) of 28 after accumulation to target value of 1e5. MS/MS acquisition was performed at a resolution of 15,000.

#### Spike-in

For the spike in-experiments, EV peptides were diluted 8 times in 10% FA. A volume of 1.5 µl of diluted EV peptide fraction was added to the same fraction of peptides from unlabeled cells. For the indicated fractions used, see Supplementary datasheet 1. Spiked-in samples were measured by MS as described above.

#### Proteomics data analysis

Raw MS files were processed in MaxQuant (version 2.6.7) using the default settings. For the SILAC samples and their controls, labels were set to 2: heavy Arg10 (+10.008269) and Lysine-D8 (+8.050214). *Homo sapiens* FASTA file used was downloaded from Uniprot on 13 January 2025. Evidence and protein group output tables were further processed in R studio (Version 2023.09.1 Build 494) with R4.4.3^28^ and the following packages: tidyverse^29^, ggvenn^30^, ggVennDiagram^31^, ComplexHeatmap^32^, pheatmap^33^, vegan^34^, and UpsetR^35^. The mass spectrometry proteomics data have been deposited to the ProteomeXchange Consortium via the PRIDE partner^36^ repository with the dataset identifier PXD070451.

#### EV*trace* confidence selection

The following steps were taken in R. Briefly, from the evidence tables of MaxQuant output rows were removed that were + for ‘potential contaminant’ or + for ‘reverse’. Next, only the peptides were kept that contained a Heavy signal intensity > 0. The peptides were grouped based on Leading razor protein and the number of rows (meaning unique peptides) grouped per sample (called ‘experiment’) were summed up. Evidence tables from separate MS measurements were merged together to one table based on Leading razor proteins. Next, additional calculations were made which included (performed per cell type): counting in how many experiments the labeled protein was detected with at least one peptide within the 3 separate experiments (cells+EVs) (*celltype*_EV_count), the sum of all labeled peptides per protein detected within all 3 experiments for the cells to which the labeled EVs were added (*celltype*_EV_true), the sum of all labeled peptides per protein detected within all 3 experiments for the cells only (*celltype*_false), and the outcome of *celltype*_EV_true - *celltype*_false (True_False). With this, the confidence criteria were applied: low ◊ True_False < 1 OR *celltype*_EV_count < 2 & *celltype*_EV_true. Medium ◊ *celltype*_EV_count = 1 & *celltype*_EV_true ≥ 2 OR True_False < 3. High ◊ True_False > 2 & *celltype*_EV_count ≥ 2 & *celltype*_EV_true ≥ 3. From this final list, proteins with the appointed confidence could be selected and merged with cleaned up protein groups tables that contained the protein intensity values. From this joined table, rankings were made based on the sum of the intensity values.

#### Enrichment analysis

High-confidence EV*trace* proteins were submitted to STRING^37^ or Metascape^38^. In both cases, the total EV proteome (pooled of n=3, 2,384 proteins) was set as background reference. For STRING the following basic settings were used: full network, evidence network edges, all interaction sources, minimum required interaction score of 0.400, and no max number of interactions. The clustering was done based on MCL clustering with the inflation parameter set on 3. In Metascape, the traced proteins of osteoclasts and osteoblasts were pooled and duplicates were removed. The same was done for the high-confidence proteins traced in the monocytes and PCa cells. Both were added in separate columns of a file that was uploaded as a multiple gene list. A custom analysis was performed, selecting GO Biological Processes, Reactome Gene Sets, KEGG Pathway, WikiPathways, DisGeNET, and Transcription Factor Targets for the enrichment analysis. The standard settings of the enrichment analysis (min overlap, P value cutoff, min enrichment) were maintained.

## Results

### Selection of EV-donor and-recipient cells for EV*trace*

To develop and provide proof of concept for our novel EV*trace* method, PC3 cells were selected as EV-donor cells. PC3 are prostate cancer (PCa) cells derived from a bone metastasis and are known to metastasize to bone *in vivo*^39^. We purified the PCa EVs using differential (ultra)centrifugation and iodixanol density gradient centrifugation in order to achieve optimal separation of EVs from non-EV contaminants. PCa cells mainly release EVs that sediment at 100,000 × *g.* The presence of common EV proteins CD63, CD9 and TSG101 in iodixanol density gradient fractions indicated that the EVs localized to the expected densities of 1.10-1.06 g/ml (Figure 1A). In these density fractions we also detected the highest numbers of EVs using high-resolution flow cytometry (Figure 1B). For the remainder of the study, we used the 100,000 × *g* pelleted EVs isolated from density fractions 1.10-1.06 g/ml. NTA measurements showed that these EVs had a size range of 60-300 nm with the peak between 110-120 nm (Figure 1C). The isolated PCa EVs were also characterized by super-resolution microscopy using dSTORM (Figure 1D-F), which showed that the majority (75%) of the EVs were positive for all three tetraspanins (CD9, CD63 and CD81), followed by CD81 and CD9 (15%), and CD81 and CD63 (5%) (Figure 1E). Here, the peak size range of the EVs was 80-120 nm (Figure 1F).

To obtain proof of concept for the EV*trace* method, we selected four different EV-recipient cell types. First, we selected osteoclasts and osteoblasts based on previous observations that these cell types can be targeted by prostate cancer EVs^19,20^. We differentiated the osteoclasts from THP1 cells as reported elsewhere^21^ and confirmed the differentiation by tartrate-resistant acid phosphatase (TRAP) staining (Supplementary Figure 1). We selected SAOS2 cells as widely-used osteoblast cell line, since this cell line shows similarity with primary human osteoblasts with regard to key osteoblastic genes^40^. In addition, we selected THP-1 cells (monocytic cells), which are well known to efficiently interact with and internalize EVs^41,42^, and the PC3 PCa cells as EV-recipient cells. To obtain a first indication that PCa EVs interacted with these EV-recipient cells, we fluorescently labeled the EVs with a lipophilic dye (PKH26). Since this dye is known to form aggregates in the same size range as EVs, we separated these potential aggregates from the stained EVs using density gradient centrifugation. In addition, we included a PKH26-stained medium sample as dye control. We observed an increase in EV interaction for all cell types over time (Figure 1G). The osteoblasts displayed the highest binding/uptake of PCa EVs, followed by the PCa cells and osteoclasts. The monocytes showed the least interaction of the four. These results confirm that the selected recipient cells interact with the PCa EVs and are suited for EV*trace* application.

### SILAC labeling of PCa EVs

The EV*trace* method relies on tracing back stable isotope-labeled EV proteins in pools of unlabeled EV-recipient cell proteins. Using LC-MS/MS, we first confirmed that PCa EV proteins could be efficiently labeled with heavy isotopes (Figure 2A). After four passages of PCa cells in the heavy isotope culture medium, 98% of the detected peptides from lysed cells contained heavy amino acids. The collected and purified EVs from these EV-donor cells consisted of 92% heavy labeled peptides. The higher percentage of unlabeled peptides in the EVs (8%) compared to the cells (2%) was further explored. Analysis on the protein level indicated that EVs contained a higher percentage (28.8%) of proteins present as both heavy labeled and unlabeled, compared to the EV-donor cells (6.3%). The percentage of proteins that remained completely unlabeled in EVs was 1%, while for cells this was 0.5%. Thus, the majority (99%) of unique proteins associated with EVs were (at least partly) heavy labeled. Almost all (98%) of the labeled EV proteins overlapped with the pool of EV proteins from cell lines reported on Vesiclepedia^43^ (Figure 2B). We also compared the protein composition of the heavy labeled EVs to unlabeled PCa EVs (Figure 2C), which showed 81.9% overlap in protein content. The correlation between the relative intensities of the 2,393 overlapping EV proteins from labeled and unlabeled EVs was 0.87. Overall, our data shows that heavy labeling of the PCa cells resulted in labeling of the far majority of EV-associated proteins without major changes to the EV composition.

**Figure 2.**
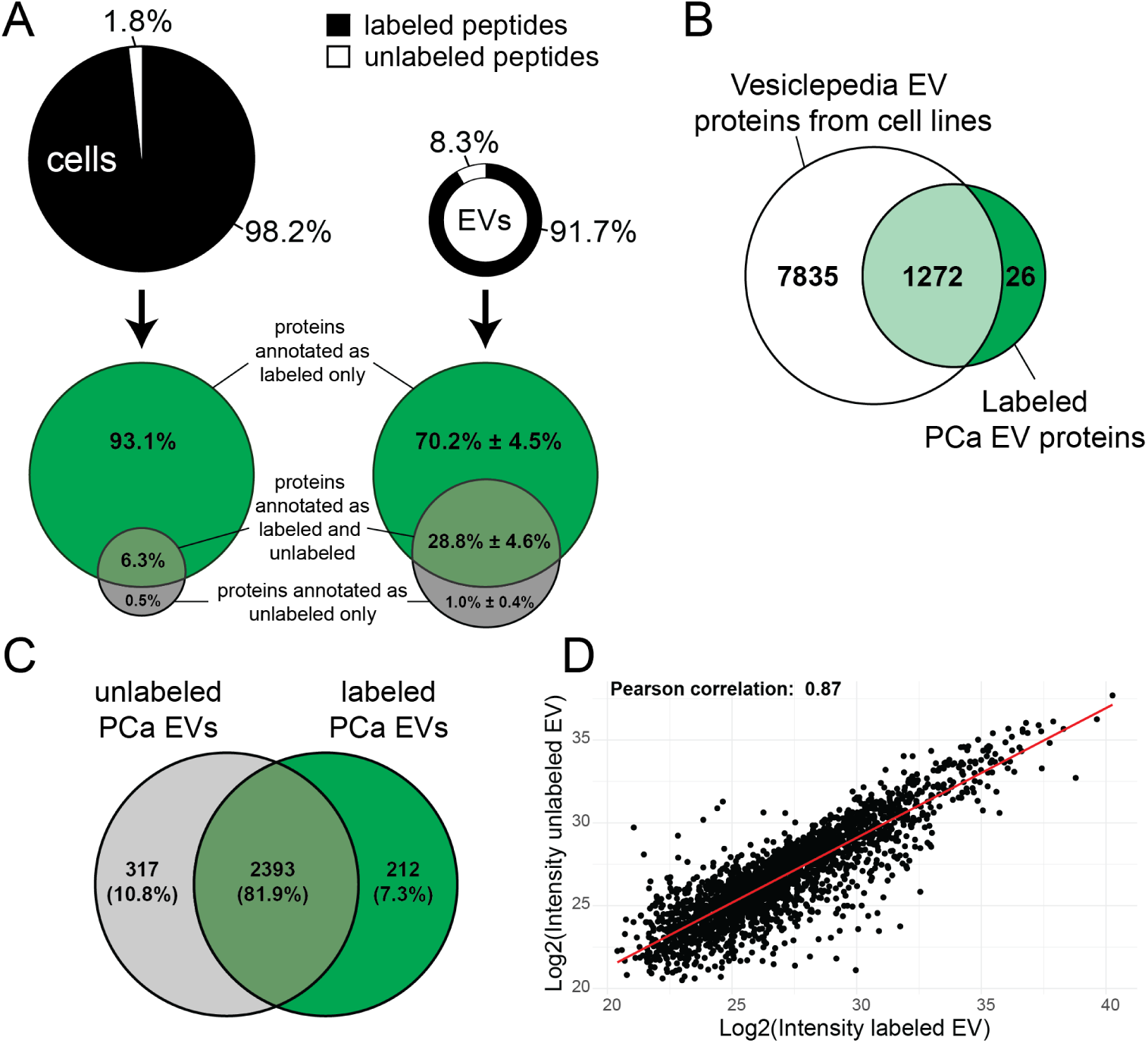
SILAC labeling of PCa EVs. (**A**) Efficiency of SILAC heavy isotope labeling of PCa cells and their released EVs. Cells were passaged for >4 times before cell harvesting and EV collection. Top: LC-MS/MS-based analysis of the percentages of the total unique peptides containing heavy amino acids (black) or not (white). Bottom: Venn diagrams of unique proteins annotated as heavy labeled (green), unlabeled (grey), or both in PCa cells and their EVs (indicated are mean % of 3 separate EV batches ± SD) (**B**) Overlap of labeled PCa EV proteins and EV proteins from various cell lines submitted to Vesiclepedia. (**C**) Venn diagram comparing the protein composition of EVs cultured in medium without (unlabeled, grey) or with (labeled, green) heavy isotopes. (**D**) Correlation plot of the overlapping 2,393 proteins in both labeled (x-axis) and unlabeled (y-axis) EVs.

### Analysis pipeline development for identification of EV*trace* proteins

Next, we incubated the heavy labeled PCa EVs with (unlabeled) osteoclasts, osteoblasts, PCa cells, and monocytes. These cell types greatly differ in size and morphology. For comparative proteomics analysis of all EV-cell incubations, we prepared cultures of each of the different EV-recipient cell type with equalized cell surface areas, as calculated by light microscopy. This resulted in more similar protein yields per cell type compared to equalizing cell numbers (Supplementary Figure 3C). This is favorable for the subsequent digestion step, which requires equal protein input. To limit the risk of EV protein breakdown after internalization, we incubated the cells with the EVs for 3 h, after which the cells were washed and lysed directly in the plate. Protein extraction and LC-MS/MS was performed on cells that were incubated with or without EVs and on the source EV population. For EV*trace*, standard SILAC quantification by heavy/light ratios of identical proteins cannot be used. Instead, low abundant heavy labeled EV proteins (EV*trace* proteins) need to be identified among high abundant pools of unlabeled cellular proteins. To obtain lists of highly confident (labeled) EV*trace* proteins, we developed a novel analysis pipeline for the annotated MS spectra.

For peptide analysis we implemented high-confidence criteria on the basis of reproducibility and on the reduction of false-positive hits. First, EV*trace* proteins were considered with high confidence if they were detected in the same cell type in at least 2 out of 3 independent experiments with a total of 3 or more (unique) peptides. For example, this would include proteins for which at least 1 heavy labeled peptide was detected in all 3 experiments, or proteins for which 1 heavy labeled peptide was detected in one experiment and a minimum of 2 unique labeled peptides in at least one other experiment. Second, we reduced the likelihood that EV*trace* hits were falsely identified as heavy labeled due to incorrect spectral matches. We assessed the probability that proteins might be incorrectly annotated as heavy labeled by analyzing the spectra of control unlabeled cells that had not been incubated with EVs. Indeed, we observed that several proteins were falsely annotated as heavy labeled peptides in these unlabeled samples (Figure 3A). A few of these proteins were falsely annotated as heavy labeled in all 4 cell types, but the majority of the false-positive hits were cell type specific (Figure 3B), suggesting that the annotation errors depended on the specific mass spectra of the sample matrices from the different cell types. To reduce false-positive EV*trace* hits, we therefore included as a high-confidence criterium that the total number of heavy labeled unique peptides of a protein detected in the EV-incubated cells needed to exceed the number of unique peptides falsely annotated as heavy labeled in the control samples (without EVs) of the same cell type by at least 3 unique peptides. A summary of these criteria for highly confident EV*trace* proteins are visualized in Figure 3C.

**Figure 3.**
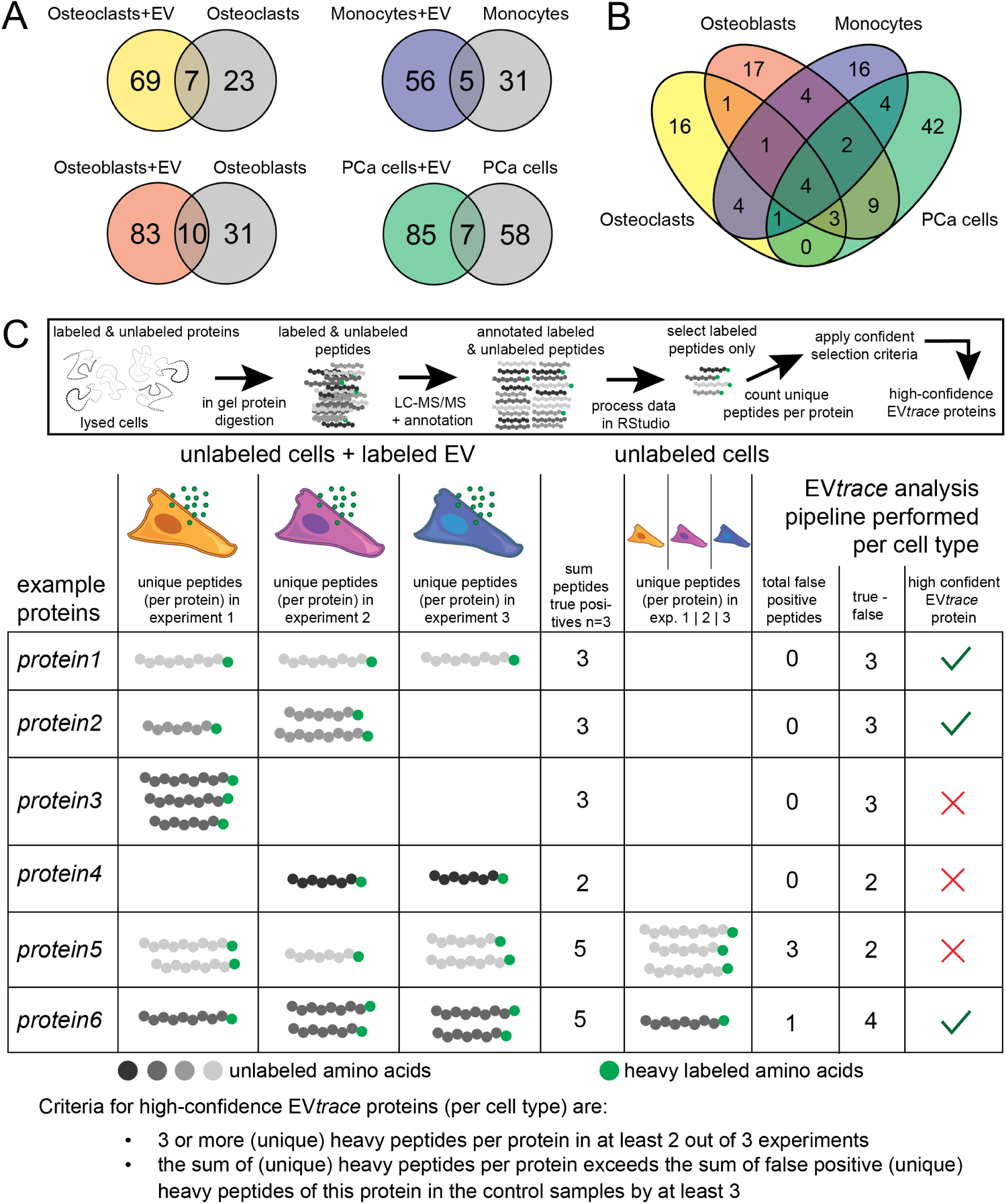
Confidence selection criteria for EV*trace* proteins. (**A**) Number of proteins annotated as heavy labeled in samples of the four different unlabeled EV-recipient cell types incubated with (color) or without (grey) heavy isotope labeled PCa EVs. Labeled proteins assigned as ‘low-confidence’ (as described in the methods section)) were excluded for the cells + EVs. For conditions of cells only, the included proteins were detected in at least 2 experiments or with >1 unique peptide. (**B**) Overlap in false-positive labeled proteins annotated in samples of the unlabeled cells only (same as grey circles in A). (**C**) Visual summary of the proteomics sample processing and analysis (top). Heavy labeled peptides were selected from all annotated peptides (MS/MS data) and unique labeled peptides per protein per sample were counted. The sum of unique labeled peptides of a protein per cell type with or without EVs was used to determine the confidence for this protein in that cell type. EV*trace* proteins were considered high-confidence when the total number of heavy labeled unique peptides of a protein detected in the EV-incubated cells exceeded the number of unique peptides falsely annotated as heavy labeled in the control samples (without EVs) by at least 3 unique peptides. Proteins in the table do not represent actual data but serve as examples showing when proteins would pass or not as highly confident following the selection criteria.

### EV*trace* protein profiles differ per recipient cell type

Using the newly developed EV*trace* data analysis pipeline, we obtained lists of high-confidence EV*trace* proteins for each of the EV-recipient cell types (Table 1). The total number of highly confident EV*trace* hits per cell type followed a similar trend as the EV uptake experiments in Figure 1E. Most high-confidence EV*trace* proteins were detected in the osteoblasts (21) and the osteoclasts (20), followed by the prostate cancer cells (17) and the monocytes (12). Interestingly, we observed little overlap in EV*trace* proteins detected in each of the different cell types (Figure 4A), even though these cells were incubated with the same pool of EVs. Of the total 49 high-confidence EV*trace* proteins, 13 were traced back in more than one cell type. Of these, only three proteins (annexin A2, integrin-β1, and histone H2B type 1) were traced back in all four cell types. The distinctive nature of the high-confidence EV*trace* protein profiles detected in the different cell types was further confirmed by hierarchical clustering (Figure 4B), which showed that EV*trace* protein profiles from each individual experiment clustered together per cell type. This clustering per EV-recipient cell type remained when lower confident traced proteins were included in the analysis (Supplementary Figure 5). The significant effect of cell type on the high-confidence traced proteins was additionally confirmed by a Pearson’s Chi-squared test (*X*-squared = 13.073, df = 3, p-value < 0.01). These results suggest that different EV-recipient cell types bind and/or internalize different subpopulations of the total pool of EVs released by PCa cells.

**Figure 4.**
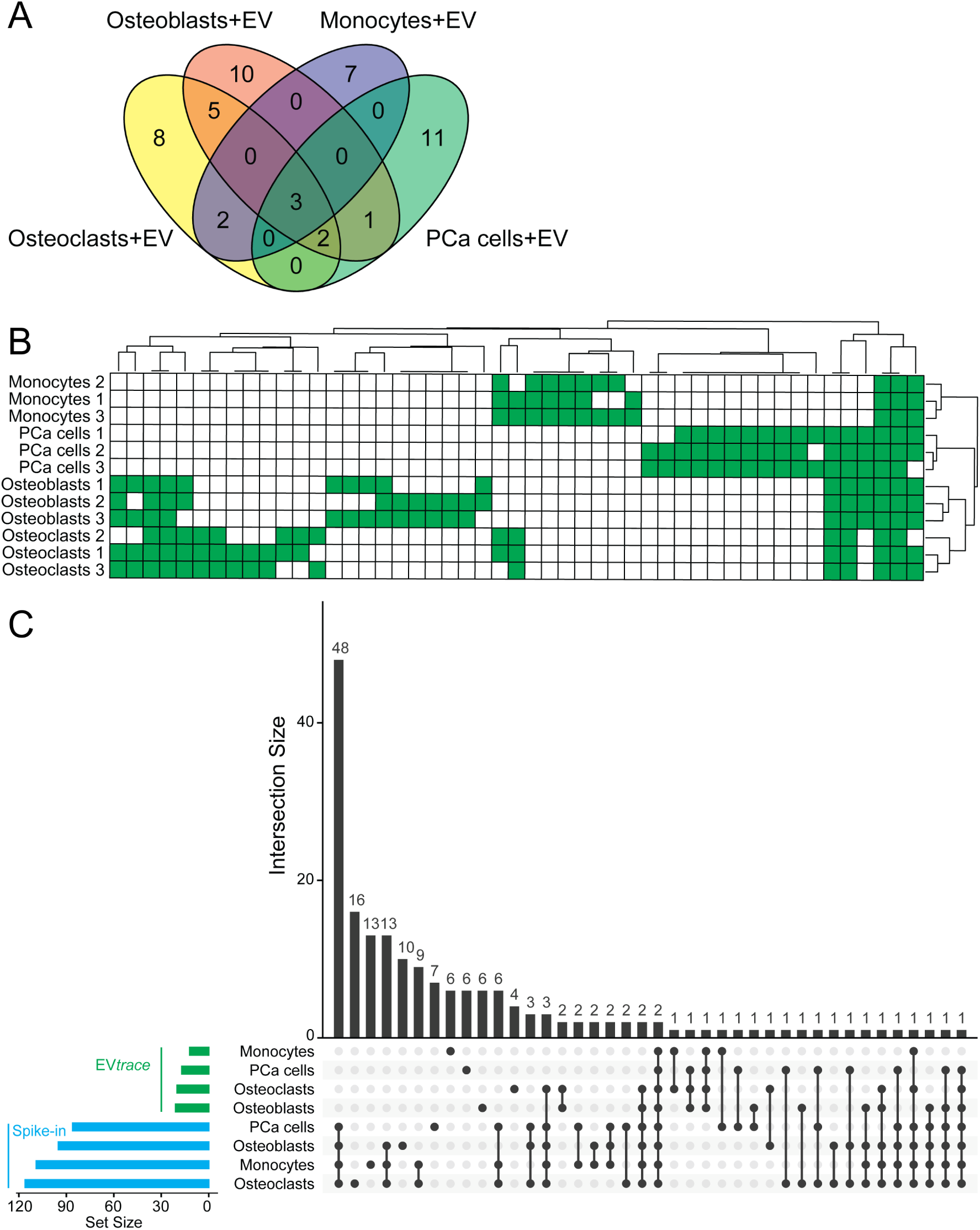
Differences in high-confidence EV*trace* protein profiles between recipient cells. (**A**) Venn diagram of the numbers and overlap of highly confident EV*trace* proteins detected in the different EV-recipient cell types. (**B**) Hierarchical clustering of the high-confidence EV*trace* proteins per EV-recipient cell type in each individual biological replicate (indicated as 1|2|3) (presence = green). Clustering was based on the Jaccard distance between the cells. (**C**) UpSet plot visualizing the overlap between high-confidence protein sets (rows) per EV-recipient cell type from EV*trace* and spiked-in EV proteins. The horizontal bars on the left-hand side indicate the total number (Set Size) of highly confident proteins in each cell type from EV*trace* (top, green) and spiked-in EV proteins (bottom, blue). On the right side, each column represents a unique combination of sets, indicated by the connected dots, and the number of shared proteins (Intersection Size) for each combination is indicated by the vertical black bars. PCa, prostate cancer

**Table 1.**
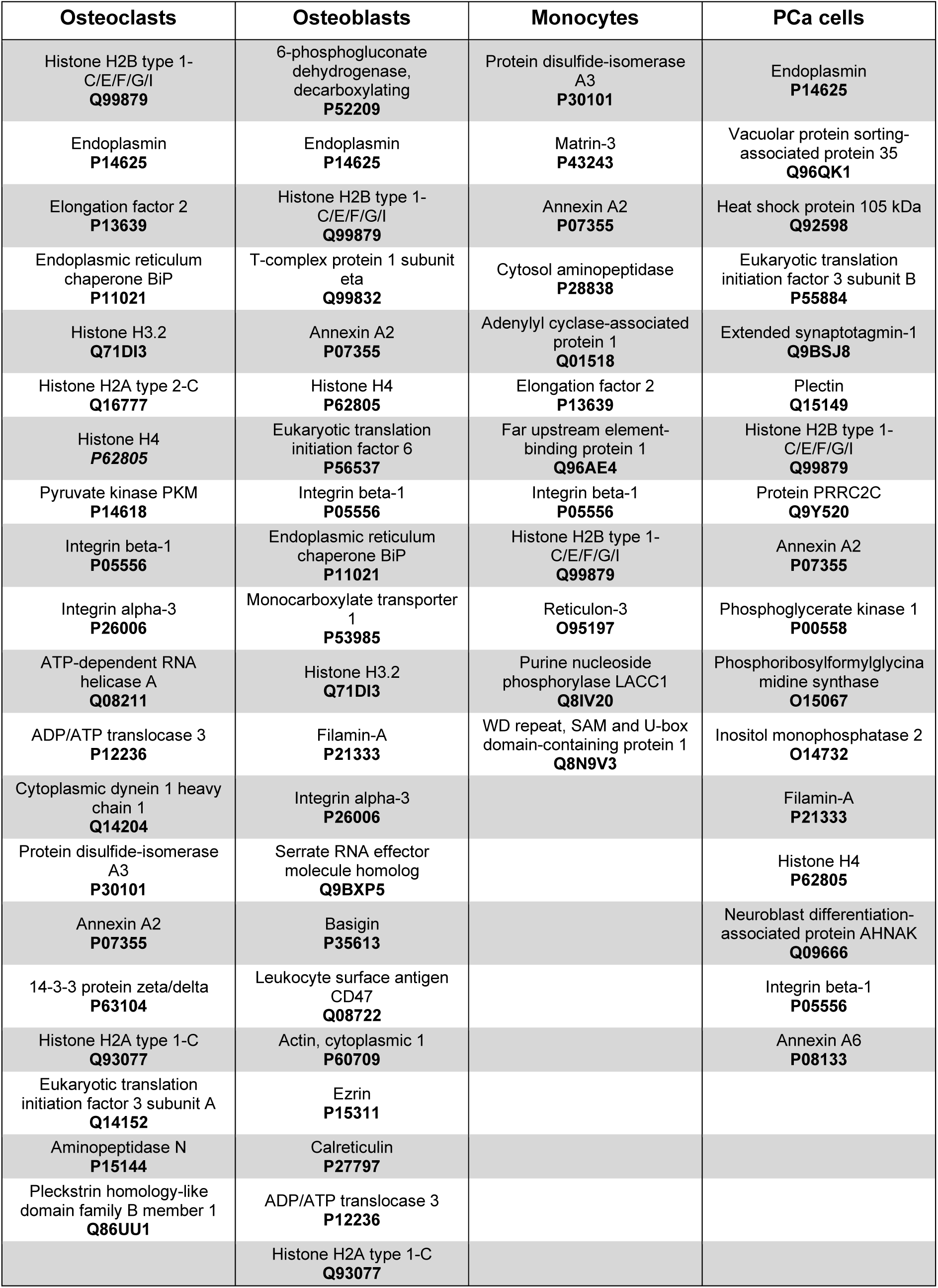
List of high-confidence traced EV proteins per cell type. Protein names and their UniProt accession number (in bold for visual separation) are listed from bottom to top as they are ranked (per cell type) by relative abundance (sum of relative intensity of n=3), with the highest ranked on top.

We next challenged the idea that specific EV subpopulations are targeted to EV-recipient cells by testing the alternative possibility that unlabeled proteins of each cell type differentially influenced which heavy labeled proteins could be traced back. In theory, it is possible that high peaks of unlabeled peptides from the cells can overshadow less abundant labeled EV peptides in a mass spectrum. To test whether the efficiency of detecting EV*trace* proteins depended on the different protein contents of the EV-recipient cells, we performed a spike-in experiment. LC-MS/MS was performed on samples in which ∼5% of digested peptides from labeled EVs was spiked into digested peptides of unlabeled cells not exposed to EVs. We applied the same criteria as in Figure 3C to identify highly confident labeled peptides. The UpSet plot showed that 27 of the 49 traced EV proteins were not detected with high confidence in the spike-in experiment, while 21 proteins identified in both the EV*trace* and the spike-in experiments were not detected in one corresponding individual cell type (Figure 4C). In addition, hierarchical clustering-based analysis of the heavy labeled proteins obtained in the spike-in experiment did not show the strict clustering per EV-recipient cell type as observed for the EV*trace* proteins (Supplementary Figure 5). These results support the conclusion that the differences in detected EV*trace* proteins between the cell types are due to specific EV-cell interactions and cannot be explained by differential interference of EV-recipient cell proteins in MS-based detection of the labeled EV proteins. Thus, the proof-of-concept experiments using our novel EV*trace* method demonstrate that different cell types internalize different sets of proteins present in the total EV pool, which are likely associated to different EV subpopulations.

### High and low abundant EV proteins are traced back in cells

The EV*trace* data described above indicate that only small fractions of the total pool of EV proteins are traced back in the EV-recipient cells with high confidence (12-21 traced proteins out of 2,378 total EV proteins). This could suggest that the sensitivity of our technology is limited and biases towards detection of highly abundant proteins, or that cells mainly interact with highly abundant EV subsets. To investigate in detail whether the EV*trace* method only traced back EV proteins that were highly abundant in the original EV protein pool, we first made a ranking of the relative abundance of the labeled PCa EV protein pool, with the highest abundant protein ranked as 1 and the lowest as 0.0004 (1/2,378). We next created similar rankings for the high-confidence EV*trace* proteins per EV-recipient cell type, and matched the proteins between the EV pool and EV*trace* (Figure 5A). The results show that many of the EV*trace* proteins in each cell type were in the top 100 most abundant proteins of the EV source, but that the ranking profiles differed between EV*trace* and EV source proteins. In addition, multiple detected EV*trace* proteins were low abundant in the EV source. For comparison, we made a similar ranking analysis of the spike-in data (Figure 5B). In contrast to the EV*trace* proteins, the vast majority of spiked-in labeled proteins were highly abundant in the source and followed a similar trend in ranking as the EV source proteins. These results indicate that EV proteins traced back in target cells are not a mere reflection of the total EV protein pool, but include selective sets of high and low abundant proteins that may co-occur on defined EV subpopulations.

**Figure 5.**
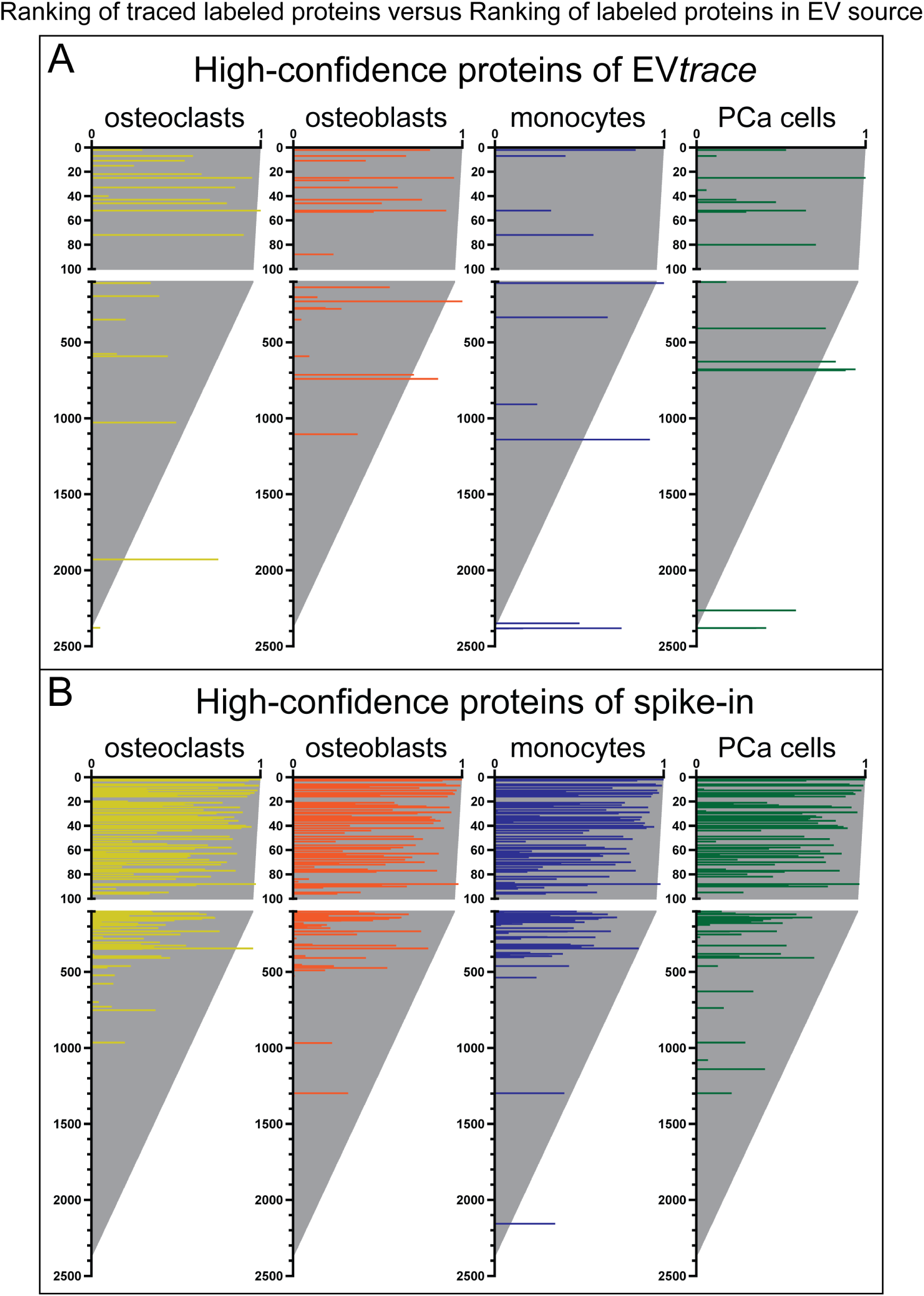
Abundance ranking of traced EV proteins. Relative abundance of labeled proteins from the PCa EV source (grey lines) were ranked from high (1) to low (1/*(total proteins)*). Superimposed are the ranking of the high-confidence EV*trace* proteins (**A**) or high-confidence spike-in proteins (**B**) for each of the EV-recipient cell type (colored lines). The same proteins of the source that are also traced back in the cells or in the spike-in samples are depicted on the same horizontal line within the ranking. The length of the colored lines indicates the relative abundance of the traced proteins.

### Identity and predicted function of traced EV proteins

It is often suggested that EVs contain sets of molecules that can collectively influence biological processes. Therefore we investigated whether the sets of traced proteins per recipient cell type were physically or functionally linked using the STRING database^37^. For assessing enrichment in protein-protein interaction (PPI), we compared EV*trace* proteins to the EV source proteome, since comparison to the full human proteome would only highlight common EV-associated networks. STRING analysis showed that the vast majority of EV*trace* protein sets detected in osteoclasts, osteoblasts, and PCa cells were significantly enriched as interaction partners (Figure 6A). For the monocytes, over half of the traced proteins were connected but there was no significant PPI enrichment, which might be due to the low number of detected proteins. Comparing the clusters of PPIs in each of the different EV-recipient cells showed little similarities (Supplementary Table 1), which suggests that the sets of EV proteins traced in each of the cell types function in different processes and pathways.

**Figure 6.**
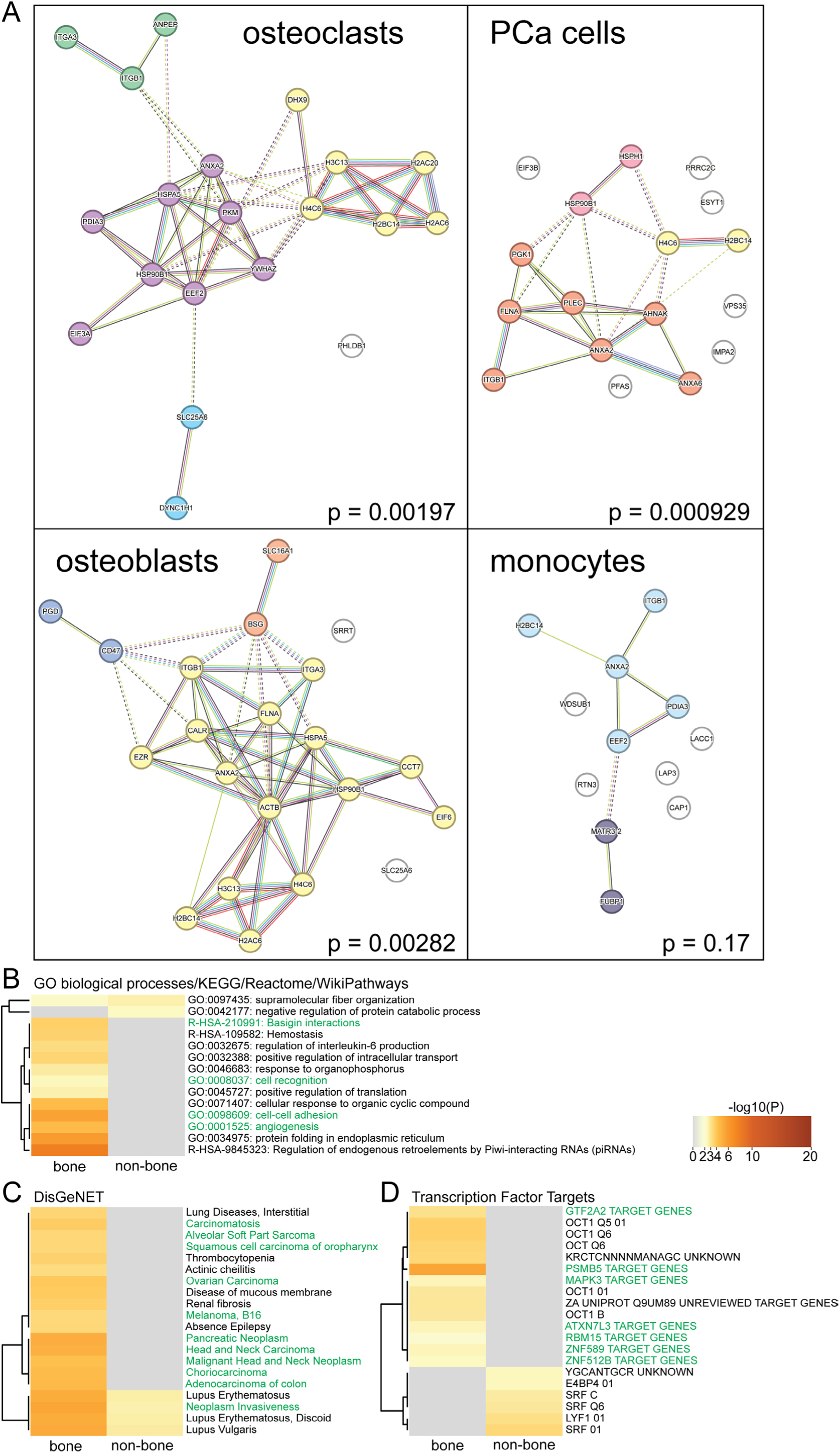
Enrichment analysis of EV*trace* proteins. (A) Network plot of high-confidence EV*trace* proteins detected in each of the different EV-recipient cells. Indicated are p-values of the protein-protein interaction (PPI) enrichment, using the EV source proteome as background. Cluster analysis was done based on standard MCL. Detailed description of cluster and protein names in Supplement table 1. (B-D) Heatmaps of significantly enriched biological processes (B, WikiPathways), diseases (C, DisGeNET), and transcription factor targets (D) for high-confidence EV*trace* proteins detected in bone (osteoclasts and osteoblasts) and non-bone (monocytes and prostate cancer (PCa) cells) cells, analyzed in Metascape. Terms associated with cancer are depicted in green.

To zoom in on potential functional pathways linked to EV proteins traced back in the different target cells, we performed an enrichment analysis in Metascape^38^ using datasets for the GO term biological processes, KEGG pathways, Reactome, and WikiPathways (Figure 6B). We compared proteins traced back in either bone cells (osteoclasts and osteoblast) or non-bone cells (PCa cells and monocytes). This allowed us to include the data from the monocytes that otherwise may not give any significant results. We noted that EV proteins traced in bone cells (but not in non-bone cells) were significantly enriched for processes important in cancer, including basigin interactions^44^, cell recognition, cell-cell adhesion, and angiogenesis. Enrichment analysis using the DisGeNET database for disease-related proteins demonstrated that EV proteins traced in the bone cells were significantly associated with various forms of cancer, neoplasm invasiveness, and carcinomatosis (Figure 6C). This set of EV proteins was also associated with transcription factor targets known to play a role in prostate cancer, such as *PSMB5*^45^ and *GTF2A2*^46^ (Figure 6D). These results support the conclusion that bone cells and non-bone cells internalized functionally different sets of EV proteins. Additionally, our data indicate that the protein profile of the prostate cancer EV subpopulation traced back in bone cells (individual proteins listed in Table 1) is congruent with a proposed role of (prostate) cancer EVs in metastasis^19,20^. Examples of cancer-related EV*trace* proteins identified in bone cells included integrin-α3, integrin-β1 and GRP78 (both bone cell types), CD47 and basigin (osteoblasts), and CD13 (osteoclasts). The proteins CD47 and integrin-α3 may be involved in binding/uptake of the EVs by bone cells. CD47, a common ‘don’t eat me’ signal when it binds to the SIRPα receptor on recipient cells^47^, was exclusively traced back in osteoblasts. By assessing the unlabeled proteomes of the recipient cells, we indeed confirmed that the SIRPα receptor was present in the osteoclasts, monocytes and PCa cells but was absent in the osteoblasts. CD47 on EVs has been shown to limit their uptake by circulating monocytes^48^ and inhibition of CD47 is being explored as anti-cancer (immuno)therapy^49^. Integrin-α3, which is known to pair with integrin-β1 on various cancer EVs, including PCa^50^, is increased on EVs in urine from PCa patients with metastasis compared to benign hyperplasia^51^. The endoplasmic reticulum chaperone BiP (also known as GRP78 or HSPA5) is a cancer-associated ER protein that is often overexpressed and mislocalized to the cell surface in various cancers^52^. GRP78 is involved in the interaction of metastatic prostate cancer cells with the bone microenvironment^53^. The protein basigin (CD147) has also been detected on EVs from other cancer origins, and is regarded as a promising drug target to reduce cancer progression^54,55^.

### Traced proteins on EV subpopulations

The differences in traced EV proteins per cell type suggests that the total pool of PCa EVs contains different EV subpopulations. To further confirm that EV*trace* proteins detected in different target cell types are present on distinct EV subpopulations, we analyzed the presence of 3 cancer-related membrane-associated proteins on individual PCa EVs using super-resolution microscopy. We selected integrin-β1 for the EV population that was taken up by all recipient cells, GRP78 for the subpopulation that targeted to the bone cells, and CD47 that was only traced back in the osteoblasts. EVs were stained with three different antibody labeling panels (Fig 7A, B) and non-specific staining was assessed using isotype control antibodies (Supplementary Figure 6A).

**Figure 7.**
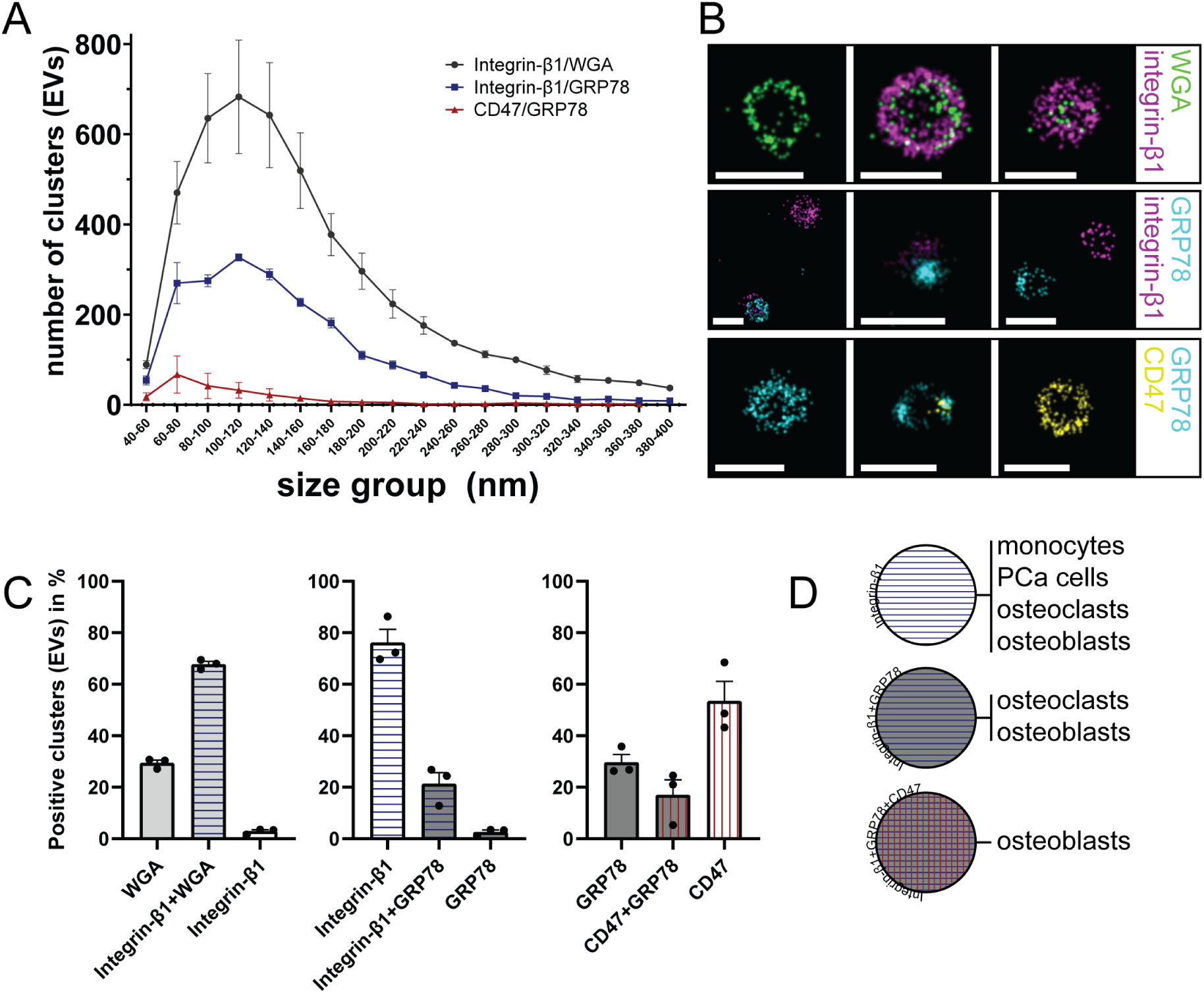
PCa EV subpopulations confirmed by super-resolution microscopy. (**A**) Distribution of the amount of clusters (indicative of EVs) per size range (nm, depicting the diameter of the clusters) detected using the different staining panels (black circle, Integrin-β1+WGA; blue square, Integrin-β1+GFRP78; red triangle, CD47+GRP78). (**B**) Representative images of individual EVs that were enriched in either one (left, right) or two (middle) of the indicated markers: WGA (green), integrin-β1 (magenta), GRP78 (cyan), CD47 (yellow). Each horizontal row represents one of the three staining panels. White scale bars indicate 200 nm. (**C**) The percentage of EVs (of all clusters detected in each staining panel as in A) that are single or double positive for indicated markers. Bars are colored to indicate markers: light grey, WGA; blue horizontal stripes, integrin-β1; dark grey, GRP78; red vertical stripes, CD47. (**D**) Graphical summary of the detected PCa EV subpopulations and in which cell types these markers were traced (see Table 1). Colors indicating the proteins are similar as in C. A and C show mean±SEM of 3 fields of view. WGA, Wheat Germ Agglutinin

EV analysis with the 3 different staining panels indicated that the majority of stained EVs were in the 80-140 nm size range (Figure 7A). These data also indicated that the quantity of detected EVs (defined as clusters with a minimum number of marker localizations) was highest when staining for more abundant proteins (e.g. integrin-β1 and WGA), and decreased with lower abundant proteins (e.g. GRP78 and CD47). EVs in the images were assessed for being single or double positive for the indicated proteins (Figure 7B, Supplementary Figure 6B), after which the number of single or double positive clusters (EVs) (Supplementary Figure 6C) and the proportion of EV subpopulations (Figure 7C) were calculated. The data indicate that the majority of WGA positive EVs also contained integrin-β1. Only part of the integrin-β1 positive EVs also contained GRP78, and the GRP78 positive population could be further divided into EVs with and without CD47. These data confirm that groups of EV proteins traced back in particular cell types can co-occur on the same EVs and mark specific EV subpopulations (Figure 7D).

Overall, the results illustrate how EV*trace* yields valuable insight into candidate proteins that can be further explored for their role in specific cell targeting and signaling by EV subpopulations in EV-donor/recipient cell pairs of interest.

## Discussion

Over the last decade, important progress has been made in the development of technologies for isolation and characterization of EVs, as well as in the analysis of their effects on cells, organs, and organisms^2^. Current technologies allow large-scale, in-depth analysis of the molecular contents of bulk EV populations using various omics-based methods^56^. In addition, high-end microscopic techniques allow assessment of EV heterogeneity by visualizing individual molecules on single EVs^8^. Yet, our knowledge on how the heterogeneous EV pool of a single cell type interacts with cells remained limited^3^. Here we presented the development and application of EV*trace*, a novel, unbiased SILAC proteomics-based method to identify EV-associated proteins that are bound or taken up by recipient cells. We demonstrated the applicability of the method by tracing proteins from prostate cancer (PCa) EVs in different recipient cell types. Using the new method we demonstrated that specific subpopulations of PCa EVs ended up in cells, and that different cell types interacted with different EV subpopulations present in the total population of PCa EVs. We validated the existence of EV populations containing the EV*trace*-identified proteins by super-resolution imaging.

A few previous studies applied SILAC as a tool to trace EV proteins, but with limited scope and potential. Some of these studies employed a reverse strategy, adding unlabeled EVs to heavy isotope-labeled cells^16–18^. An important downside of this approach is that insufficiently labeled proteins in EV-recipient cells could be mistakenly interpreted as internalized unlabeled EV proteins. The same holds true for unlabeled FBS proteins that contaminate EV preparations, which could be wrongly annotated as human (EV) proteins due to conserved sequences^57^. Moreover, previous studies often depended on ratiometric analysis of labeled/unlabeled (e.g. heavy/light) proteins, or focused on tracking back pre-defined proteins^15,16^. This precluded an unbiased analysis of the EV fractions that were preferentially internalized by target cells.

### Technical considerations relevant for the EV*trace* method

During the development of EV*trace*, we identified key factors for appropriate application of the method and reliable identification of internalized EV subpopulations.

#### Isotope labeling conditions

Results obtained with EV*trace* are most robust when using highly purified EVs and high-level heavy labeling of EV proteins. Yet, trace contaminations with FBS EVs or incomplete labeling of EV proteins do not compromise data validity, as EV*trace* analysis only selects labeled EV peptides and proteins. The primary limitation of lower labeling efficiency of EV proteins is a reduced probability of detection and high-confidence traceability.

#### Optimizing EV-to-target cell ratios

It is not recommended to standardize EV-to-recipient cell ratios by cell number if recipient cells vary substantially in size and morphology. This can introduce bias in the EV*trace* procedure due to unequal protein input, which affects downstream proteomics. To minimize this risk, it is best to equalize surface area coverage per condition (Supplementary Figure 3). Furthermore, we advise for each condition to consider the balance between uptake efficiency and protein degradation and to optimize EV concentration and incubation time. Given that EV sources are often limited, the well size and culture volume can be reduced to maximize the EV-to-cell ratio. We here report tracing of EV proteins after a 3 hour incubation period. Longer incubation times allow for increased EV internalization (Figure 1), but also raise the likelihood of lysosomal routing and degradation of EV proteins. Whether a longer incubation time increases or decreases the number of traced EV proteins and affects the interpretation of EV*trace* data remains to be elucidated.

#### Proper controls and their importance

It is crucial for EV*trace* to include a sample of recipient cells not incubated with EVs. Our data clearly demonstrate that low-level detection of false-positive hits (proteins incorrectly annotated as heavy labeled in unlabeled cells) cannot be prevented, and that the identity of false-positive hits varies per cell type (Figure 3B). Inclusion of unlabeled conditions of each recipient cell types is therefore crucial to correct for these false-positives.

A second important control sample is the total EV population that is used to add to recipient cells. Assessing the proteome of this ‘source’ EV population is crucial to identify and assess (selective) uptake of traced EV subpopulations (Figure 5).

### Application of EV*trace* to progress the EV field

EV*trace* can be used to answer fundamental questions in the field of EV biology regarding the (selective) uptake of EV subpopulations. It can generate valuable sources of candidate proteins that can be further explored for their role in EV binding/uptake and functional modification of target cells. To showcase this, we applied EV*trace* to identify PCa EV subpopulations that are preferentially taken up by bone cells, a common metastatic site in PCa^19^. Our results demonstrated strong coherence between the traced EV proteins targeting bone cells and protein clusters and pathways involved in cancer progression and metastasis (Figure 6). While some traced proteins are known players in metastasis and therapeutic targets^51,53–55^, many have not yet been associated with cell targeting or tumor development, suggesting that EV*trace* could uncover novel candidates for therapeutic intervention.

Beyond cancer research, EV*trace* offers broad applications for studying EV protein transfer between cells in diverse biological contexts without the need for cell engineering. The primary requirement is that EV-donor cells undergo sufficient divisions for efficient isotope labeling. Alternatively, pulse labeling could be explored in systems with enhanced EV biogenesis/release, though this may require validation to ensure labeling levels are detectable and traceable. EV*trace* can also be employed to compare EV (sub)populations and their cellular targeting across different (engineered) EV-donor cell types or under varying culture conditions, to identify features of the source EV proteome that influence targeting efficiency and/or specificity. Besides *in vitro* application, as exemplified in our current study, EV*trace* also holds promise for *in vivo* applications. Although obtaining sufficient labeled material poses a challenge, SILAC-labeled EVs have been successfully isolated from *in vivo* models^15,58,59^, opening opportunities to investigate how EVs from various body fluids interact with recipient cells in physiological contexts.

While our focus has been on tracing EV-associated proteins and identifying (sub)populations internalized by EV-recipient cells, EV*trace* simultaneously captures the full proteome of the unlabeled recipient cells. This enables exploration of receptor-ligand interactions mediating EV-cell communication and assessment of proteomic changes induced by EV uptake.

Together, these possibilities highlight EV*trace* as a versatile approach with substantial potential for advancing our understanding of EV biology and intercellular communication in health and disease.

## Acknowledgements

We are grateful to the Centre for Cellular Imaging Utrecht and the Flow Cytometry and Cell Sorting Facility, both at the Faculty for Veterinary Medicine of Utrecht University, for allowing access to their facilities. The authors would like to thank Dr. Lorenzo Albertazzi and Stijn van Veen for their feedback on the super-resolution microscopy data and all partners of the Horizon 2020 FET Open MimicKeY consortium for their helpful feedback during this project.

