## supplemental data file for "EV*trace*: tracing extracellular vesicles-associated proteins in recipient cells using stable isotope labeling"

| Experiment 1 | Gel fraction | name of raw file |
|---|---|---|
| OC+EV1 | 1.1 | 20230106_EX1_UM8_Damen101_SA_ext02_MEK_1_1 |
| OC+EV1 | 1.2 | 20221219_EX1_UM8_Damen101_SA_ext02_MEK_1_2 |
| OC+EV1 | 1.3 | 20221219_EX1_UM8_Damen101_SA_ext02_MEK_1_3 |
| OC+EV1 | 1.4 | 20221219_EX1_UM8_Damen101_SA_ext02_MEK_1_4 |
| OC+EV1 | 1.5 | 20221219_EX1_UM8_Damen101_SA_ext02_MEK_1_5 |
| OC+EV1 | 1.6 | 20221219_EX1_UM8_Damen101_SA_ext02_MEK_1_6 |
| OC+EV1 | 1.7 | 20221219_EX1_UM8_Damen101_SA_ext02_MEK_1_7 |
| OC+EV1 | 1.8 | 20221219_EX1_UM8_Damen101_SA_ext02_MEK_1_8 |
| OC+EV1 | 1.9 | 20221219_EX1_UM8_Damen101_SA_ext02_MEK_1_9 |
| OC 1 | 2.1 | 20221220_EX1_UM8_Damen101_SA_ext02_MEK_2_1 |
| OC 1 | 2.2 | 20221220_EX1_UM8_Damen101_SA_ext02_MEK_2_2 |
| OC 1 | 2.3 | 20221220_EX1_UM8_Damen101_SA_ext02_MEK_2_3 |
| OC 1 | 2.4 | 20221220_EX1_UM8_Damen101_SA_ext02_MEK_2_4 |
| OC 1 | 2.5 | 20221220_EX1_UM8_Damen101_SA_ext02_MEK_2_5 |
| OC 1 | 2.6 | 20221220_EX1_UM8_Damen101_SA_ext02_MEK_2_6 |
| OC 1 | 2.7 | 20221220_EX1_UM8_Damen101_SA_ext02_MEK_2_7 |
| OC 1 | 2.8 | 20221220_EX1_UM8_Damen101_SA_ext02_MEK_2_8 |
| OC 1 | 2.9 | 20221220_EX1_UM8_Damen101_SA_ext02_MEK_2_9 |
| PC3+EV1 | 3.1 | 20221220_EX1_UM8_Damen101_SA_ext02_MEK_3_1 |
| PC3+EV1 | 3.2 | 20221220_EX1_UM8_Damen101_SA_ext02_MEK_3_2 |
| PC3+EV1 | 3.3 | 20221220_EX1_UM8_Damen101_SA_ext02_MEK_3_3 |
| PC3+EV1 | 3.4 | 20221220_EX1_UM8_Damen101_SA_ext02_MEK_3_4 |
| PC3+EV1 | 3.5 | 20221220_EX1_UM8_Damen101_SA_ext02_MEK_3_5 |
| PC3+EV1 | 3.6 | 20221220_EX1_UM8_Damen101_SA_ext02_MEK_3_6 |
| PC3+EV1 | 3.7 | 20221220_EX1_UM8_Damen101_SA_ext02_MEK_3_7 |
| PC3+EV1 | 3.8 | 20221220_EX1_UM8_Damen101_SA_ext02_MEK_3_8 |
| PC3+EV1 | 3.9 | 20221220_EX1_UM8_Damen101_SA_ext02_MEK_3_9 |
| PC3 1 | 4.1 | 20221221_EX1_UM8_Damen101_SA_ext02_MEK_4_1 |
| PC3 1 | 4.2 | 20221221_EX1_UM8_Damen101_SA_ext02_MEK_4_2 |
| PC3 1 | 4.3 | 20221221_EX1_UM8_Damen101_SA_ext02_MEK_4_3 |
| PC3 1 | 4.4 | 20221221_EX1_UM8_Damen101_SA_ext02_MEK_4_4 |
| PC3 1 | 4.5 | 20221221_EX1_UM8_Damen101_SA_ext02_MEK_4_5 |
| PC3 1 | 4.6 | 20221221_EX1_UM8_Damen101_SA_ext02_MEK_4_6 |
| PC3 1 | 4.7 | 20221221_EX1_UM8_Damen101_SA_ext02_MEK_4_7 |
| PC3 1 | 4.8 | 20221221_EX1_UM8_Damen101_SA_ext02_MEK_4_8 |
| PC3 1 | 4.9 | 20221221_EX1_UM8_Damen101_SA_ext02_MEK_4_9 |
| SOAS2+EV1 | 5.1 | 20221221_EX1_UM8_Damen101_SA_ext02_MEK_5_1 |
| SOAS2+EV1 | 5.2 | 20221221_EX1_UM8_Damen101_SA_ext02_MEK_5_2 |
| SOAS2+EV1 | 5.3 | 20221221_EX1_UM8_Damen101_SA_ext02_MEK_5_3 |
| SOAS2+EV1 | 5.4 | 20221221_EX1_UM8_Damen101_SA_ext02_MEK_5_4 |
| SOAS2+EV1 | 5.5 | 20230106_EX1_UM8_Damen101_SA_ext02_MEK_5_5 |
| SOAS2+EV1 | 5.6 | 20221221_EX1_UM8_Damen101_SA_ext02_MEK_5_6 |
| SOAS2+EV1 | 5.7 | 20221221_EX1_UM8_Damen101_SA_ext02_MEK_5_7 |
| SOAS2+EV1 | 5.8 | 20221221_EX1_UM8_Damen101_SA_ext02_MEK_5_8 |
| SOAS2+EV1 | 5.9 | 20221221_EX1_UM8_Damen101_SA_ext02_MEK_5_9 |
| SAOS2 1 | 6.1 | 20221222_EX1_UM8_Damen101_SA_ext02_MEK_6_1 |
| SAOS2 1 | 6.2 | 20221222_EX1_UM8_Damen101_SA_ext02_MEK_6_2 |
| SAOS2 1 | 6.3 | 20221222_EX1_UM8_Damen101_SA_ext02_MEK_6_3 |
| SAOS2 1 | 6.4 | 20221222_EX1_UM8_Damen101_SA_ext02_MEK_6_4 |
| SAOS2 1 | 6.5 | 20221222_EX1_UM8_Damen101_SA_ext02_MEK_6_5 |
| SAOS2 1 | 6.6 | 20221222_EX1_UM8_Damen101_SA_ext02_MEK_6_6 |
| SAOS2 1 | 6.7 | 20221222_EX1_UM8_Damen101_SA_ext02_MEK_6_7 |
| SAOS2 1 | 6.8 | 20221222_EX1_UM8_Damen101_SA_ext02_MEK_6_8 |
| SAOS2 1 | 6.9 | 20221222_EX1_UM8_Damen101_SA_ext02_MEK_6_9 |
| THP1+EV1 | 7.1 | 20221222_EX1_UM8_Damen101_SA_ext02_MEK_7_1 |
| THP1+EV1 | 7.2 | 20221222_EX1_UM8_Damen101_SA_ext02_MEK_7_2 |
| THP1+EV1 | 7.3 | 20230106_EX1_UM8_Damen101_SA_ext02_MEK_7_3 |
| THP1+EV1 | 7.4 | 20221222_EX1_UM8_Damen101_SA_ext02_MEK_7_4 |
| THP1+EV1 | 7.5 | 20221222_EX1_UM8_Damen101_SA_ext02_MEK_7_5 |
| THP1+EV1 | 7.6 | 20221222_EX1_UM8_Damen101_SA_ext02_MEK_7_6 |
| THP1+EV1 | 7.7 | 20221222_EX1_UM8_Damen101_SA_ext02_MEK_7_7 |
| THP1+EV1 | 7.8 | 20221222_EX1_UM8_Damen101_SA_ext02_MEK_7_8 |
| THP1+EV1 | 7.9 | 20221222_EX1_UM8_Damen101_SA_ext02_MEK_7_9 |
| THP1 1 | 8.1 | 20221223_EX1_UM8_Damen101_SA_ext02_MEK_8_1 |
| THP1 1 | 8.2 | 20221223_EX1_UM8_Damen101_SA_ext02_MEK_8_2 |
| THP1 1 | 8.3 | 20221223_EX1_UM8_Damen101_SA_ext02_MEK_8_3 |
| THP1 1 | 8.4 | 20221223_EX1_UM8_Damen101_SA_ext02_MEK_8_4 |
| THP1 1 | 8.5 | 20221223_EX1_UM8_Damen101_SA_ext02_MEK_8_5 |
| THP1 1 | 8.6 | 20230106_EX1_UM8_Damen101_SA_ext02_MEK_8_6 |
| THP1 1 | 8.7 | 20221223_EX1_UM8_Damen101_SA_ext02_MEK_8_7 |
| THP1 1 | 8.8 | 20221223_EX1_UM8_Damen101_SA_ext02_MEK_8_8 |
| THP1 1 | 8.9 | 20221223_EX1_UM8_Damen101_SA_ext02_MEK_8_9 |
| EV1 | 11.1 | 20221223_EX1_UM8_Damen101_SA_ext02_MEK_11_1 |
| EV1 | 11.2 | 20221223_EX1_UM8_Damen101_SA_ext02_MEK_11_2 |
| EV1 | 11.3 | 20221223_EX1_UM8_Damen101_SA_ext02_MEK_11_3 |
| EV1 | 11.4 | 20221223_EX1_UM8_Damen101_SA_ext02_MEK_11_4 |
| EV1 | 11.5 | 20221223_EX1_UM8_Damen101_SA_ext02_MEK_11_5 |
| EV1 | 11.6 | 20221223_EX1_UM8_Damen101_SA_ext02_MEK_11_6 |
| EV1 | 11.7 | 20221223_EX1_UM8_Damen101_SA_ext02_MEK_11_7 |
| EV1 | 11.8 | 20221223_EX1_UM8_Damen101_SA_ext02_MEK_11_8 |
| EV1 | 11.9 | 20221223_EX1_UM8_Damen101_SA_ext02_MEK_11_9 |

| Experiment 2 | Gel fraction | Fraction nr after pool | name of raw file |
|---|---|---|---|
| OC+EV2 | 1.1 | 1 | 20230414_EX3_UM7_damen101_SA_EXT02_MEK_1 |
| OC+EV2 | 1.2 | | |
| OC+EV2 | 1.3 | 2 | 20230414_EX3_UM7_damen101_SA_EXT02_MEK_2 |
| OC+EV2 | 1.4 | | |
| OC+EV2 | 1.5 | 3 | 20230414_EX3_UM7_damen101_SA_EXT02_MEK_3 |
| OC+EV2 | 1.6 | 4 | 20230414_EX3_UM7_damen101_SA_EXT02_MEK_4 |
| OC+EV2 | 1.7 | 5 | 20230414_EX3_UM7_damen101_SA_EXT02_MEK_5 |
| OC+EV2 | 1.8 | 6 | 20230414_EX3_UM7_damen101_SA_EXT02_MEK_6 |
| OC+EV2 | 1.9 | | |
| OC+EV3 | 2.1 | 7 | 20230414_EX3_UM7_damen101_SA_EXT02_MEK_7 |
| OC+EV3 | 2.2 | | |
| OC+EV3 | 2.3 | 8 | 20230414_EX3_UM7_damen101_SA_EXT02_MEK_8 |
| OC+EV3 | 2.4 | | |
| OC+EV3 | 2.5 | 9 | 20230414_EX3_UM7_damen101_SA_EXT02_MEK_9 |
| OC+EV3 | 2.6 | 10 | 20230414_EX3_UM7_damen101_SA_EXT02_MEK_10 |
| OC+EV3 | 2.7 | 11 | 20230414_EX3_UM7_damen101_SA_EXT02_MEK_11 |
| OC+EV3 | 2.8 | 12 | 20230414_EX3_UM7_damen101_SA_EXT02_MEK_12 |
| OC+EV3 | 2.9 | | |
| OC+EV3 | 3.1 | 13 | 20230414_EX3_UM7_damen101_SA_EXT02_MEK_13 |
| OC 2 | 3.2 | | |
| OC 2 | 3.3 | 14 | 20230414_EX3_UM7_damen101_SA_EXT02_MEK_14 |
| OC 2 | 3.4 | | |
| OC 2 | 3.5 | 15 | 20230414_EX3_UM7_damen101_SA_EXT02_MEK_15 |
| OC 2 | 3.6 | 16 | 20230414_EX3_UM7_damen101_SA_EXT02_MEK_16 |
| OC 2 | 3.7 | 17 | 20230414_EX3_UM7_damen101_SA_EXT02_MEK_17 |
| OC 2 | 3.8 | 18 | 20230414_EX3_UM7_damen101_SA_EXT02_MEK_18 |
| OC 2 | 3.9 | | |
| PC3+EV2 | 4.1 | 19 | 20230414_EX3_UM7_damen101_SA_EXT02_MEK_19 |
| PC3+EV2 | 4.2 | | |
| PC3+EV2 | 4.3 | 20 | 20230414_EX3_UM7_damen101_SA_EXT02_MEK_20 |
| PC3+EV2 | 4.4 | | |
| PC3+EV2 | 4.5 | 21 | 20230414_EX3_UM7_damen101_SA_EXT02_MEK_21 |
| PC3+EV2 | 4.6 | 22 | 20230414_EX3_UM7_damen101_SA_EXT02_MEK_22 |
| PC3+EV2 | 4.7 | 23 | 20230414_EX3_UM7_damen101_SA_EXT02_MEK_23 |
| PC3+EV2 | 4.8 | 24 | 20230414_EX3_UM7_damen101_SA_EXT02_MEK_24 |
| PC3+EV2 | 4.9 | | |
| PC3+EV3 | 5.1 | 25 | 20230414_EX3_UM7_damen101_SA_EXT02_MEK_25 |
| PC3+EV3 | 5.2 | | |
| PC3+EV3 | 5.3 | 26 | 20230414_EX3_UM7_damen101_SA_EXT02_MEK_26 |
| PC3+EV3 | 5.4 | | |
| PC3+EV3 | 5.5 | 27 | 20230414_EX3_UM7_damen101_SA_EXT02_MEK_27 |
| PC3+EV3 | 5.6 | 28 | 20230414_EX3_UM7_damen101_SA_EXT02_MEK_28 |
| PC3+EV3 | 5.7 | 29 | 20230414_EX3_UM7_damen101_SA_EXT02_MEK_29 |
| PC3+EV3 | 5.8 | 30 | 20230414_EX3_UM7_damen101_SA_EXT02_MEK_30 |
| PC3+EV3 | 5.9 | | |
| PC3 2 | 6.1 | 31 | 20230417_EX3_UM7_damen101_SA_EXT02_MEK_31 |
| PC3 2 | 6.2 | | |
| PC3 2 | 6.3 | 32 | 20230417_EX3_UM7_damen101_SA_EXT02_MEK_32 |
| PC3 2 | 6.4 | | |
| PC3 2 | 6.5 | 33 | 20230417_EX3_UM7_damen101_SA_EXT02_MEK_33 |
| PC3 2 | 6.6 | 34 | 20230417_EX3_UM7_damen101_SA_EXT02_MEK_34 |
| PC3 2 | 6.7 | 35 | 20230417_EX3_UM7_damen101_SA_EXT02_MEK_35 |
| PC3 2 | 6.8 | 36 | 20230417_EX3_UM7_damen101_SA_EXT02_MEK_36 |
| PC3 2 | 6.9 | | |
| SOAS2+EV2 | 7.1 | 37 | 20230417_EX3_UM7_damen101_SA_EXT02_MEK_37 |
| SOAS2+EV2 | 7.2 | | |
| SOAS2+EV2 | 7.3 | 38 | 20230417_EX3_UM7_damen101_SA_EXT02_MEK_38 |
| SOAS2+EV2 | 7.4 | | |
| SOAS2+EV2 | 7.5 | 39 | 20230417_EX3_UM7_damen101_SA_EXT02_MEK_39 |
| SOAS2+EV2 | 7.6 | 40 | 20230417_EX3_UM7_damen101_SA_EXT02_MEK_40 |
| SOAS2+EV2 | 7.7 | 41 | 20230417_EX3_UM7_damen101_SA_EXT02_MEK_41 |
| SOAS2+EV2 | 7.8 | 42 | 20230417_EX3_UM7_damen101_SA_EXT02_MEK_42 |
| SOAS2+EV2 | 7.9 | | |
| SAOS2+EV3 | 8.1 | 43 | 20230417_EX3_UM7_damen101_SA_EXT02_MEK_43 |
| SAOS2+EV3 | 8.2 | | |
| SAOS2+EV3 | 8.3 | 44 | 20230417_EX3_UM7_damen101_SA_EXT02_MEK_44 |
| SAOS2+EV3 | 8.4 | | |
| SAOS2+EV3 | 8.5 | 45 | 20230417_EX3_UM7_damen101_SA_EXT02_MEK_45 |
| SAOS2+EV3 | 8.6 | 46 | 20230417_EX3_UM7_damen101_SA_EXT02_MEK_46 |
| SAOS2+EV3 | 8.7 | 47 | 20230417_EX3_UM7_damen101_SA_EXT02_MEK_47 |
| SAOS2+EV3 | 8.8 | 48 | 20230417_EX3_UM7_damen101_SA_EXT02_MEK_48 |
| SAOS2+EV3 | 8.9 | | |
| SAOS2 2 | 9.1 | 49 | 20230417_EX3_UM7_damen101_SA_EXT02_MEK_49 |
| SAOS2 2 | 9.2 | | |
| SAOS2 2 | 9.3 | 50 | 20230417_EX3_UM7_damen101_SA_EXT02_MEK_50 |
| SAOS2 2 | 9.4 | | |
| SAOS2 2 | 9.5 | 51 | 20230417_EX3_UM7_damen101_SA_EXT02_MEK_51 |
| SAOS2 2 | 9.6 | 52 | 20230417_EX3_UM7_damen101_SA_EXT02_MEK_52 |
| SAOS2 2 | 9.7 | 53 | 20230417_EX3_UM7_damen101_SA_EXT02_MEK_53 |
| SAOS2 2 | 9.8 | 54 | 20230417_EX3_UM7_damen101_SA_EXT02_MEK_54 |
| SAOS2 2 | 9.9 | | |
| THP1+EV2 | 10.1 | 55 | 20230417_EX3_UM7_damen101_SA_EXT02_MEK_55 |
| THP1+EV2 | 10.2 | | |
| THP1+EV2 | 10.3 | 56 | 20230417_EX3_UM7_damen101_SA_EXT02_MEK_56 |
| THP1+EV2 | 10.4 | | |
| THP1+EV2 | 10.5 | 57 | 20230417_EX3_UM7_damen101_SA_EXT02_MEK_57 |
| THP1+EV2 | 10.6 | 58 | 20230417_EX3_UM7_damen101_SA_EXT02_MEK_58 |
| THP1+EV2 | 10.7 | 59 | 20230417_EX3_UM7_damen101_SA_EXT02_MEK_59 |
| THP1+EV2 | 10.8 | 60 | 20230417_EX3_UM7_damen101_SA_EXT02_MEK_60 |
| THP1+EV2 | 10.9 | | |
| THP1+EV3 | 11.1 | 61 | 20230417_EX3_UM7_damen101_SA_EXT02_MEK_61 |
| THP1+EV3 | 11.2 | | |
| THP1+EV3 | 11.3 | 62 | 20230417_EX3_UM7_damen101_SA_EXT02_MEK_62 |
| THP1+EV3 | 11.4 | | |
| THP1+EV3 | 11.5 | 63 | 20230417_EX3_UM7_damen101_SA_EXT02_MEK_63 |
| THP1+EV3 | 11.6 | 64 | 20230417_EX3_UM7_damen101_SA_EXT02_MEK_64 |
| THP1+EV3 | 11.7 | 65 | 20230417_EX3_UM7_damen101_SA_EXT02_MEK_65 |
| THP1+EV3 | 11.8 | 66 | 20230417_EX3_UM7_damen101_SA_EXT02_MEK_66 |
| THP1+EV3 | 11.9 | | |
| THP1 2 | 12.1 | 67 | 20230417_EX3_UM7_damen101_SA_EXT02_MEK_67 |
| THP1 2 | 12.2 | | |
| THP1 2 | 12.3 | 68 | 20230417_EX3_UM7_damen101_SA_EXT02_MEK_68 |
| THP1 2 | 12.4 | | |
| THP1 2 | 12.5 | 69 | 20230417_EX3_UM7_damen101_SA_EXT02_MEK_69 |
| THP1 2 | 12.6 | 70 | 20230417_EX3_UM7_damen101_SA_EXT02_MEK_70 |
| THP1 2 | 12.7 | 71 | 20230417_EX3_UM7_damen101_SA_EXT02_MEK_71 |
| THP1 2 | 12.8 | 72 | 20230417_EX3_UM7_damen101_SA_EXT02_MEK_72 |
| THP1 2 | 12.9 | | |
| EV2 | 16.1 | 16.1 | 20230505_EX3_UM7_Damen101_SA_EXT02_MEK_16_1 |
| EV2 | 16.2 | 16.2 | 20230505_EX3_UM7_Damen101_SA_EXT02_MEK_16_2 |
| EV2 | 16.3 | 16.3 | 20230505_EX3_UM7_Damen101_SA_EXT02_MEK_16_3 |
| EV2 | 16.4 | 16.4 | 20230505_EX3_UM7_Damen101_SA_EXT02_MEK_16_4 |
| EV2 | 16.5 | 16.5 | 20230505_EX3_UM7_Damen101_SA_EXT02_MEK_16_5 |
| EV2 | 16.6 | 16.6 | 20230505_EX3_UM7_Damen101_SA_EXT02_MEK_16_6 |
| EV2 | 16.7 | 16.7 | 20230505_EX3_UM7_Damen101_SA_EXT02_MEK_16_7 |
| EV2 | 16.8 | 16.8 | 20230505_EX3_UM7_Damen101_SA_EXT02_MEK_16_8 |
| EV2 | 16.9 | 16.9 | 20230505_EX3_UM7_Damen101_SA_EXT02_MEK_16_9 |
| EV3 | 17.1 | 17.1 | 20230505_EX3_UM7_Damen101_SA_EXT02_MEK_17_1 |
| EV3 | 17.2 | 17.2 | 20230505_EX3_UM7_Damen101_SA_EXT02_MEK_17_2 |
| EV3 | 17.3 | 17.3 | 20230505_EX3_UM7_Damen101_SA_EXT02_MEK_17_3 |
| EV3 | 17.4 | 17.4 | 20230505_EX3_UM7_Damen101_SA_EXT02_MEK_17_4 |
| EV3 | 17.5 | 17.5 | 20230505_EX3_UM7_Damen101_SA_EXT02_MEK_17_5 |
| EV3 | 17.6 | 17.6 | 20230505_EX3_UM7_Damen101_SA_EXT02_MEK_17_6 |
| EV3 | 17.7 | 17.7 | 20230505_EX3_UM7_Damen101_SA_EXT02_MEK_17_7 |
| EV3 | 17.8 | 17.8 | 20230505_EX3_UM7_Damen101_SA_EXT02_MEK_17_8 |
| EV3 | 17.9 | 17.9 | 20230505_EX3_UM7_Damen101_SA_EXT02_MEK_17_9 |

Spike-in

1,5 µL 1:8 labeled EV fraction
+ 1,5 µL unlabeld cell fraction

| EV fraction | cell fraction | Spiked-in sample nr | name of raw file |
|---|---|---|---|
| 11.4 | 2.4 | 1 | 20230718_EX1_UM8_Damen101_SA_EXT0_MEK_sp1 |
| | 4.4 | 2 | 20230718_EX1_UM8_Damen101_SA_EXT0_MEK_sp2 |
| | 6.4 | 3 | 20230718_EX1_UM8_Damen101_SA_EXT0_MEK_sp3 |
| | 8.4 | 4 | 20230718_EX1_UM8_Damen101_SA_EXT0_MEK_sp4 |
| 11.7 | 2.7 | 5 | 20230718_EX1_UM8_Damen101_SA_EXT0_MEK_sp5 |
| | 4.7 | 6 | 20230718_EX1_UM8_Damen101_SA_EXT0_MEK_sp6 |
| | 6.7 | 7 | 20230718_EX1_UM8_Damen101_SA_EXT0_MEK_sp7 |
| | 8.7 | 8 | 20230718_EX1_UM8_Damen101_SA_EXT0_MEK_sp8 |
| 16.3+16.4 | 14 | 9 | 20230718_EX1_UM8_Damen101_SA_EXT0_MEK_sp9 |
| | 32 | 10 | 20230718_EX1_UM8_Damen101_SA_EXT0_MEK_sp10 |
| | 50 | 11 | 20230718_EX1_UM8_Damen101_SA_EXT0_MEK_sp11 |
| | 68 | 12 | 20230718_EX1_UM8_Damen101_SA_EXT0_MEK_sp12 |
| 16.7 | 17 | 13 | 20230718_EX1_UM8_Damen101_SA_EXT0_MEK_sp13 |
| | 35 | 14 | 20230718_EX1_UM8_Damen101_SA_EXT0_MEK_sp14 |
| | 53 | 15 | 20230718_EX1_UM8_Damen101_SA_EXT0_MEK_sp15 |
| | 71 | 16 | 20230718_EX1_UM8_Damen101_SA_EXT0_MEK_sp16 |
| 17.3+17.4 | 14 | 17 | 20230718_EX1_UM8_Damen101_SA_EXT0_MEK_sp17 |
| | 32 | 18 | 20230718_EX1_UM8_Damen101_SA_EXT0_MEK_sp18 |
| | 50 | 19 | 20230718_EX1_UM8_Damen101_SA_EXT0_MEK_sp19 |
| | 68 | 20 | 20230718_EX1_UM8_Damen101_SA_EXT0_MEK_sp20 |
| 17.7 | 17 | 21 | 20230718_EX1_UM8_Damen101_SA_EXT0_MEK_sp21 |
| | 35 | 22 | 20230718_EX1_UM8_Damen101_SA_EXT0_MEK_sp22 |
| | 53 | 23 | 20230718_EX1_UM8_Damen101_SA_EXT0_MEK_sp23 |
| | 71 | 24 | 20230718_EX1_UM8_Damen101_SA_EXT0_MEK_sp24 |
