## supplemental information for "EV*trace*: tracing extracellular vesicles-associated proteins in recipient cells using stable isotope labeling"

### Supplementary Information

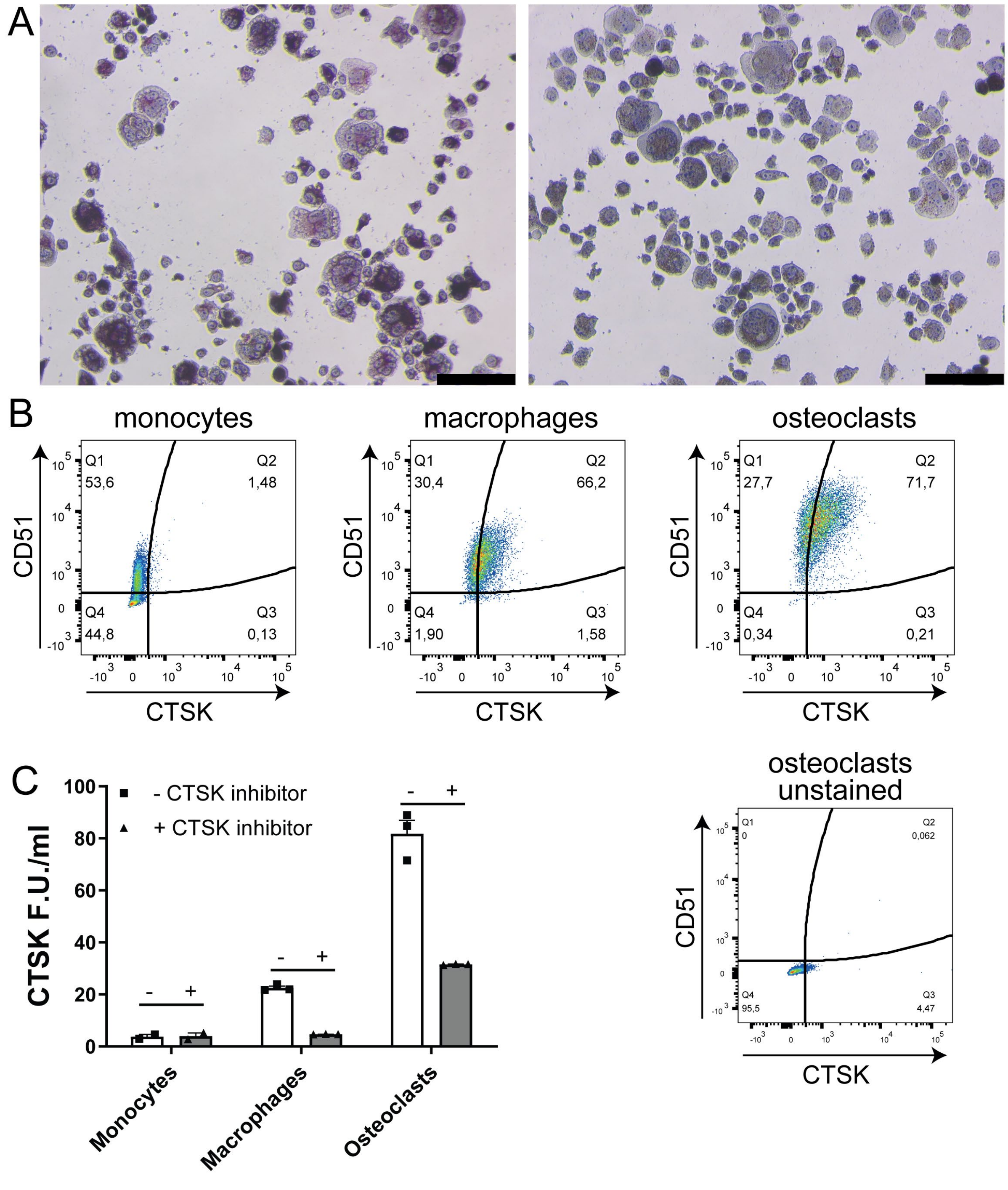


**Supplementary Figure 1.** (**A**) Tartrate resistant acid phosphatase (TRAP) staining of osteoclasts from differentiated THP1 cells. Left is acid phosphatase (purple/red), right is the tartrate resistant acid phosphatase (red/brown). Cells additionally show a multinuclear phenotype. Black scale bar is 100 µm. (**B**) Osteoclasts markers CD51 and CTSK in monocytes (THP1 cells), macrophages (PMA stimulated THP1 cells), and osteoclasts (differentiated from THP1 cells) measured by flow cytometry. (**C**) CTSK enzyme activity assay (F.U./ml) by the same cell conditions as in B. Substrate was added in the absence (squares, white bars) or presence (triangles, grey bars) of a CTSK inhibitor to confirm cleavage selectivity. CTSK; cathepsin K

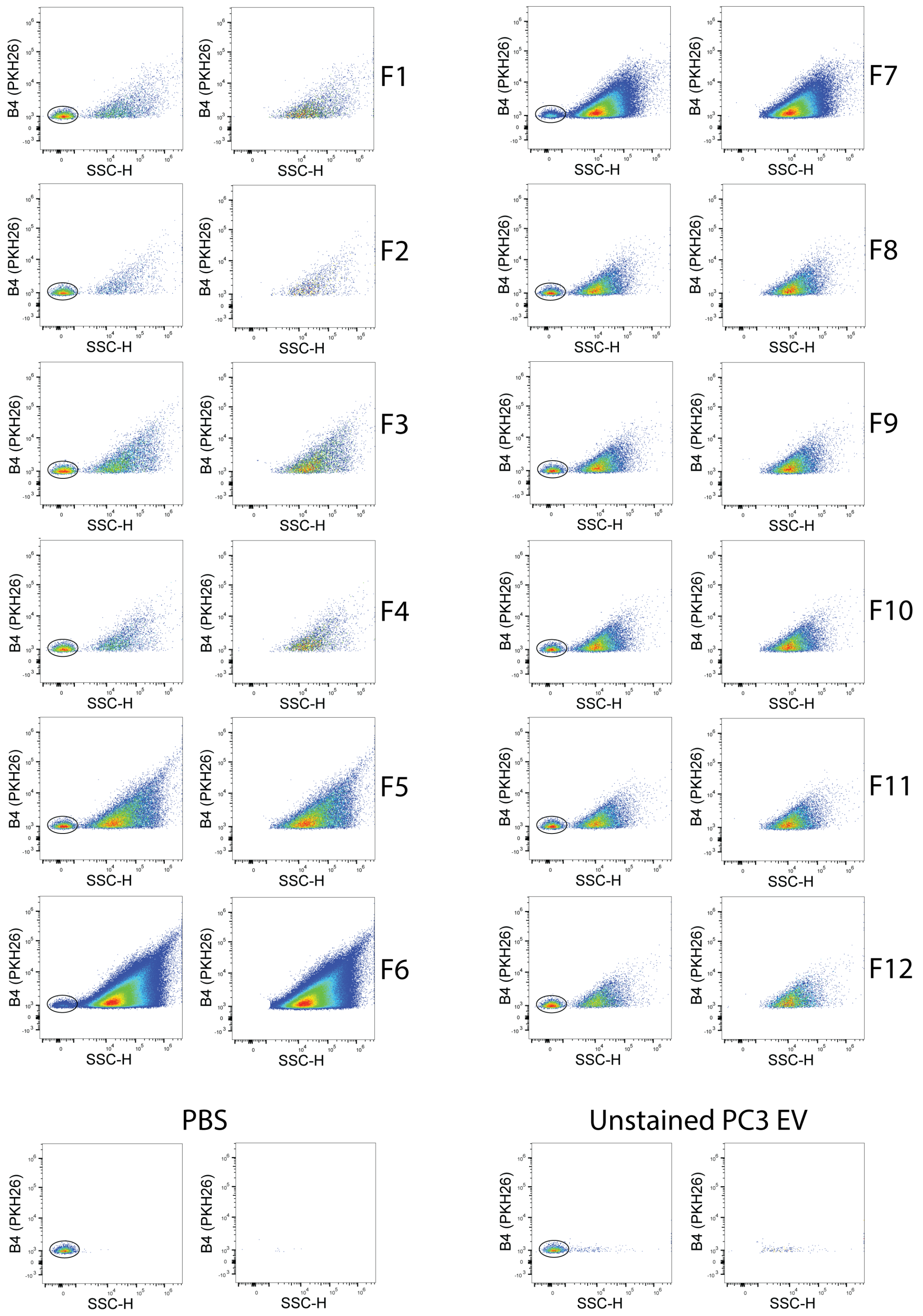


**Supplementary Figure 2.** High-resolution flow cytometric analysis of EVs detected in PKH26 stained 100,000 × *g* pellets of PCa-derived EVs separated by iodixanol density gradient centrifugation. Presented are dot plots of particles detected in fractions 1 to 12. PBS control and unstained EV samples (bottom) were included to determine the fluorescent threshold on the B4 detector. PCa, prostate cancer

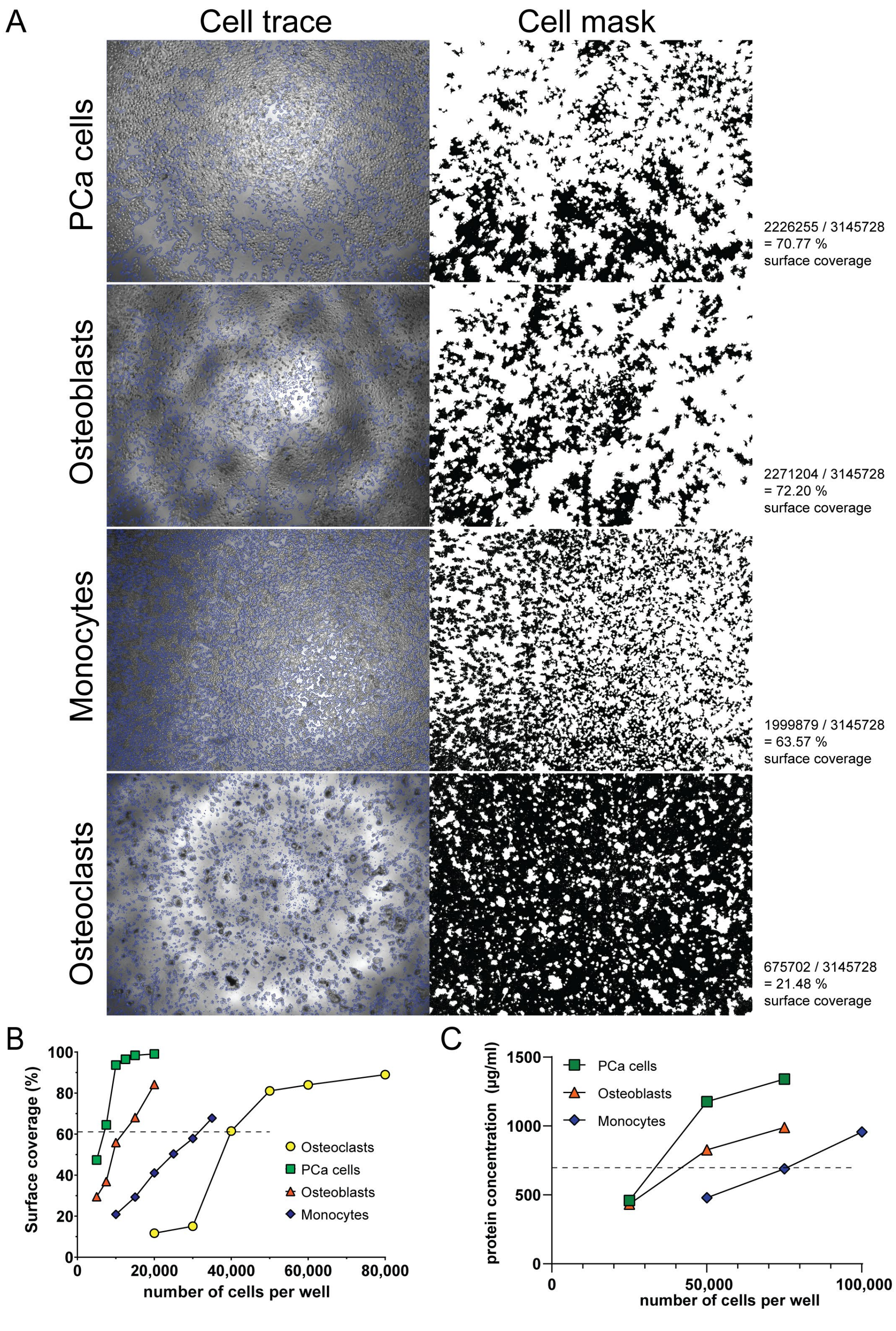


**Supplementary Figure 3.** Cell surface coverage in cultures of the 4 different EV-recipient cell types and corresponding cell and protein concentrations. (**A**) Image examples (captured with light microscopy with 4x objective) of each cell type and cell surface computation using the phase contrast microscopy segmentation toolbox PHANTAST (cell contours indicated in white). The amount of white pixels were divided by the total pixels of the image (3145728), which resulted in the percentage of surface coverage of cells in that well/image. The percentage surface coverage was used as normalization for volume of EV pool added. (**B**) Cell surface coverage (%) of each cell type per number of cells seeded in a 96 well plate. Data illustrate that different numbers of each cell type is needed to obtain similar cell coverage values. (**C**) Protein concentration (determined by BCA) of all lysed cells from one well in a 48 well plate using different cell concentrations. Data illustrate that different number of cells need to be seeded for each cell type to obtain a similar protein concentration.

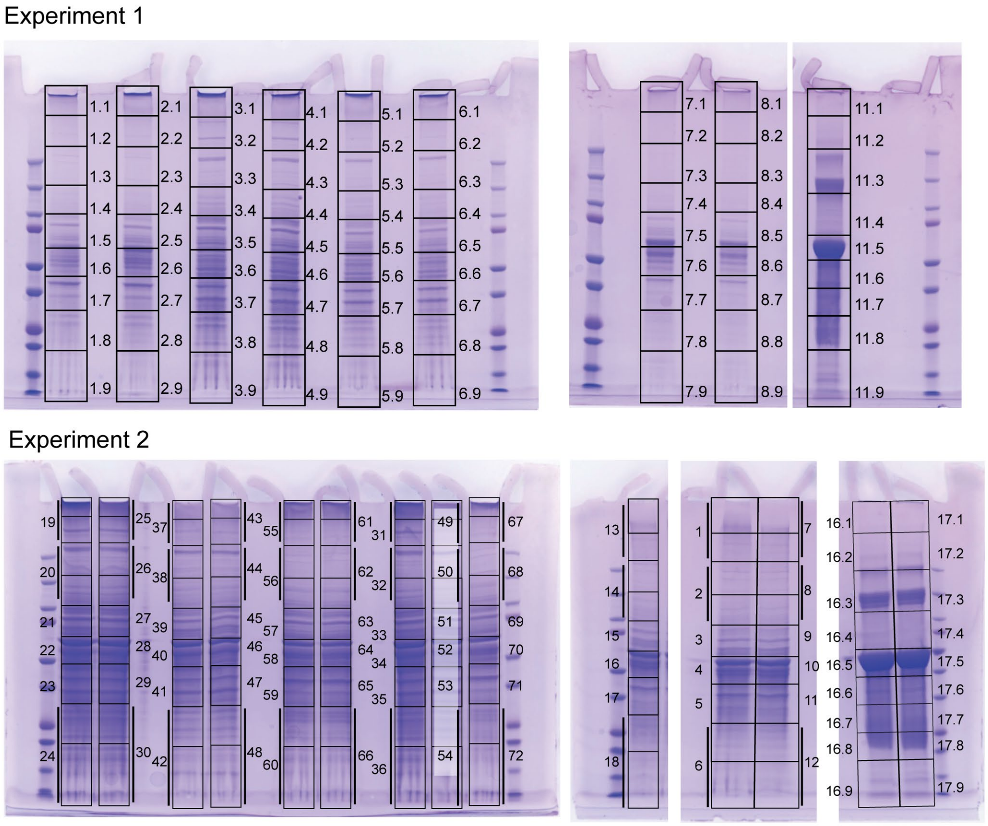


**Supplementary Figure 4.** Gel fractionations for in gel trypsin digestion of samples used for proteomics. Numbers correspond to the numbering that can be found in the names of the raw files (Supplementary Datasheet 1). Note that some fractions were pooled after digestion (indicated by solid vertical lines flanking the lanes).

**Supplementary table 1.** Explanation of EV*trace* protein clusters depicted in Figure 6A.

| | Node color | gene count | primary description | secondary description | tertiary description | protein names |
|---|---|---|---|---|---|---|
| **osteoclasts** | purple | 8 | Protein folding in endoplasmic reticulum | Endoplasmic reticulum chaperone complex | - | HSP90B1, HSPA5, ANXA2, YWHAZ, EIF3A, PKM, EEF2, PDIA3 |
| | yellow | 6 | RNA Polymerase I Promoter Opening | Systemic lupus erythematosus | Nucleosome core | H4C6, DHX9, H2AC6, H2BC14, H2AC20, H3C13 |
| | green | 3 | Integrin alpha3-beta1 complex | - | - | ANPEP, ITGB1, ITGA3 |
| | light blue | 2 | - | - | - | SLC25A6, DYNC1H1 |
| **osteoblasts** | yellow | 15 | B-WICH complex positively regulates rRNA expression | Nucleosome core | - | H4C6, HSP90B1, HSPA5, ANXA2, ACTB, H2AC6, H2BC14, H3C13, CCT7, EIF6, FLNA, CALR, ITGA3, ITGB1, EZR |
| | dark blue | 2 | - | - | - | PGD, CD47 |
| | orange | 2 | Defective SLC16A1 causes symptomatic deficiency in lactate transport (SDLT) | Proton-coupled monocarboxylate transport | - | BSG, SLC16A1 |
| **PCa cells** | red | 7 | Myelin sheath | Mixed, incl. Positive regulation of plasma membrane repair, and Annexin | - | ANXA2, AHNAK, PGK1, FLNA, PLEC, ITGB1, ANXA6 |
| | yellow | 2 | RNA Polymerase I Promoter Opening | - | - | H4C6, H2BC14 |
| | pink | 2 | Endocytic vesicle lumen | - | - | HSPH1, HSP90B1 |
| **monocytes** | light blue | 5 | - | - | - | PDIA3, ANXA2, EEF2, H2BC14, ITGB1 |
| | dark blue | 2 | - | - | - | MATR3-2, FUBP1 |

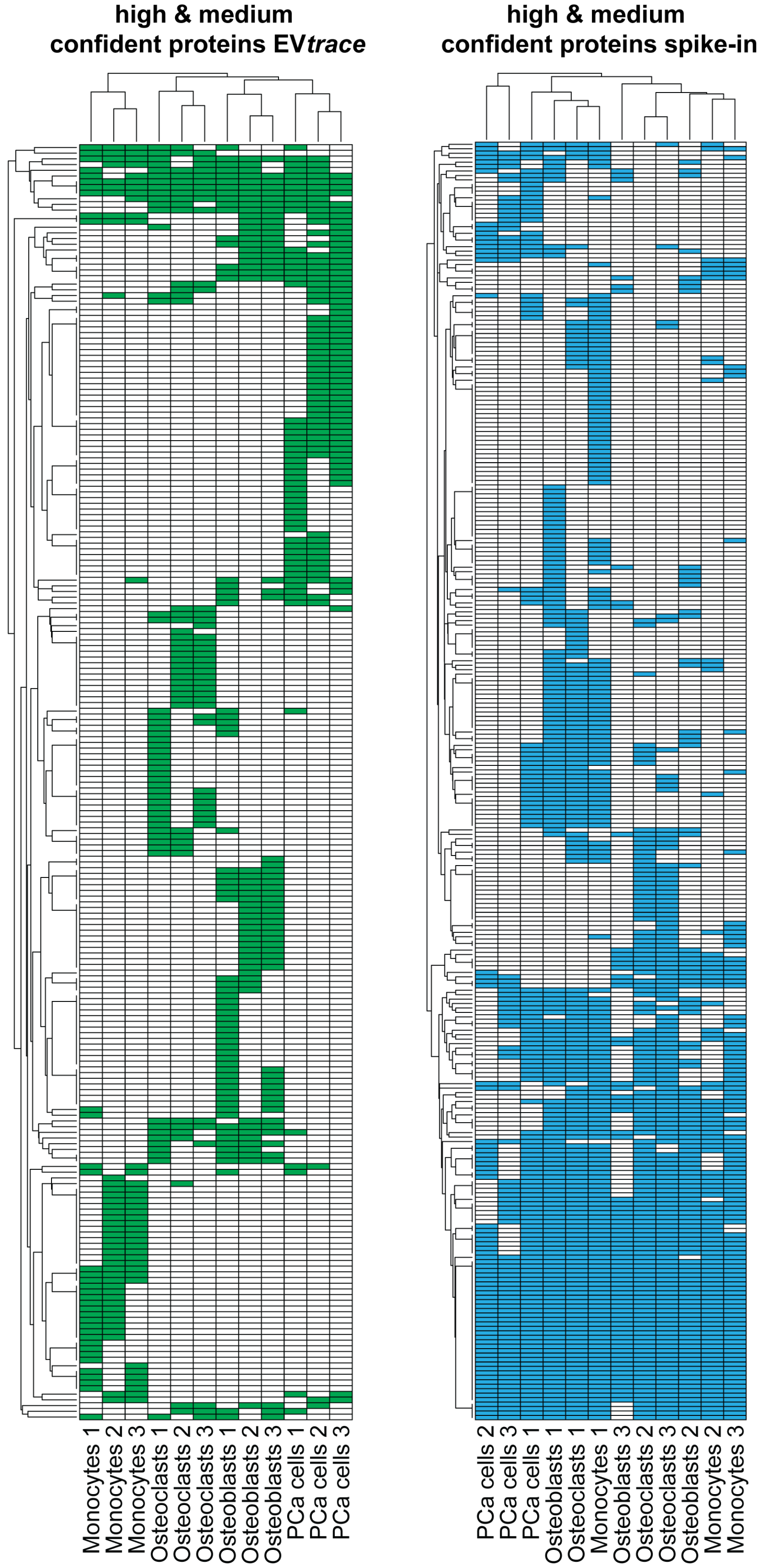


**Supplementary Figure 5.** Hierarchical clustering of high- and medium-confidence proteins of EV*trace* (left) and the spike-in (right) experiments (1-3). Clustering was based on the Jaccard distance of the EV*trace* (green) and spike in (blue) protein profiles between the cells. Data illustrate that EV*trace* proteins strictly cluster by EV-recipient cell type, whereas spike-in proteins do not.

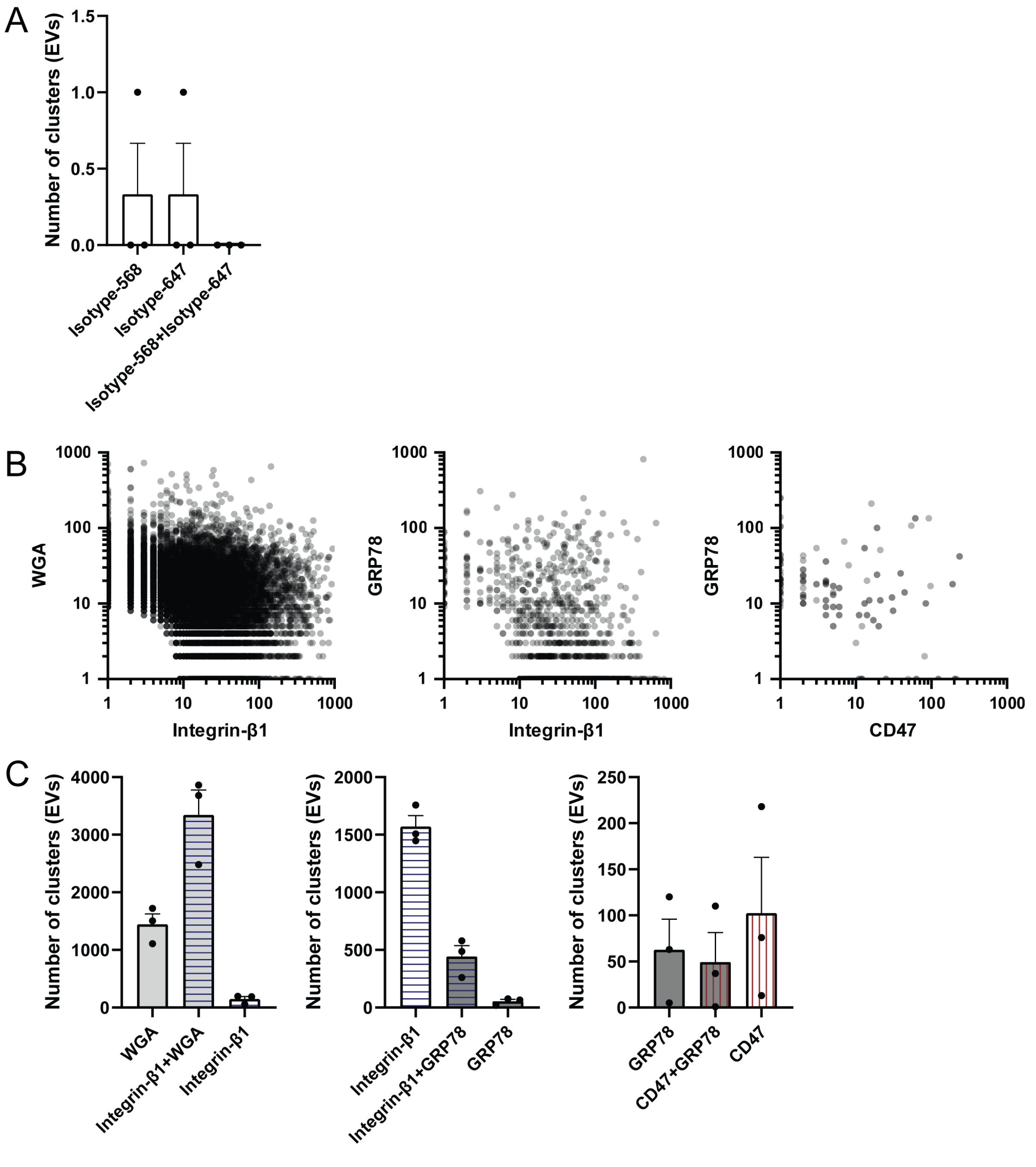


**Supplementary Figure 6.** Super-resolution microscopy of PCa EV shows subpopulations containing GRP78 with or without CD47. (**A**) PCa EVs incubated with isotype (mouse IgG1) controls were analyzed with the same settings as the other staining panels. Each point of the total number of clusters per field of view. Each cluster is indicative of an EV. (**B**) Dotplots of the double positive EVs, showing the number of localizations (X and Y axis) of the indicated markers per EV/cluster. (**C**) Number of clusters (EVs) positive for either one or both indicated proteins per staining panel, corresponding to Figure 7C. Each point represents the total number of clusters per field of view. Bar graphs in A and C show mean±SEM. WGA, Wheat Germ Agglutinin
